# Prime-pull immunization of mice with a BcfA-adjuvanted vaccine elicits mucosal immunity and prevents SARS CoV-2 infection and pathology

**DOI:** 10.1101/2022.04.06.487394

**Authors:** Mohamed M. Shamseldin, Ashley Zani, Adam Kenney, Jack Evans, Cong Zeng, Kaitlin A. Read, Kyle Caution, Jesse M. Hall, Jessica M. Brown, Gilian Gunsch, Kara N. Corps, Supranee Chaiwatpongsakorn, KC Mahesh, Mijia Lu, Rajendar Deora, Mark E. Peeples, Jianrong Li, Kenneth J. Oestreich, Shan-Lu Liu, Jacob S. Yount, Purnima Dubey

**Author notes:** equal contribution (Ashley Zani, Adam Kenney, Jack Evans, Cong Zeng, Kaitlin A. Read).

## Abstract

Vaccines against SARS-CoV-2 that induce mucosal immunity capable of preventing infection and disease remain urgently needed. We show that intramuscular priming of mice with an alum and BcfA-adjuvanted Spike subunit vaccine, followed by a BcfA-adjuvanted mucosal booster, generated Th17 polarized tissue resident CD4+ T cells, and mucosal and serum antibodies. The serum antibodies efficiently neutralized SARS-CoV-2 and its Delta variant, suggesting cross-protection against a recent variant of concern (VOC). Immunization with this heterologous vaccine prevented weight loss following challenge with mouse-adapted SARS-CoV-2 and reduced viral replication in the nose and lungs. Histopathology showed a strong leukocyte and polymorphonuclear (PMN) cell infiltrate without epithelial damage in mice immunized with BcfA-containing vaccines. In contrast, viral load was not reduced in the upper respiratory tract of IL-17 knockout mice immunized with the same formulation, suggesting that the Th17 polarized T cell responses are critical for protection. We show that vaccines adjuvanted with alum and BcfA, delivered through a heterologous prime-pull regimen, protect against SARS-CoV-2 infection without causing enhanced respiratory disease.

**SIGNIFICANCE:** There remains a need for SARS CoV-2 booster vaccines that generate mucosal immunity and prevent transmission. We show that systemic priming followed by a mucosal booster with a BcfA-adjuvanted subunit vaccine generates neutralizing antibodies and Th17 polarized systemic and tissue-resident immune responses that provide sterilizing immunity against wildtype SARS CoV-2, and a variant of concern. Importantly, in contrast to alum alone, the addition of BcfA prevents respiratory pathology. These results suggest that a BcfA-adjuvanted mucosal booster may elicit mucosal immunity in individuals previously immunized systemically with approved vaccines. This foundational study in mice sets the stage for testing our vaccine regimen in larger animal models as a booster vaccine.

## INTRODUCTION

COVID-19 is a respiratory and multi-organ disease caused by severe acute respiratory syndrome coronavirus 2 (SARS-CoV-2), a positive-sense RNA virus belonging to the family *Coronaviridae*, and the etiologic agent of the ongoing pandemic (Hsieh et al., 2020; Kaneko et al., 2020; Zhu et al., 2020). To date, more than 435 million cases of infection and nearly 6 million deaths have been reported globally making COVID-19 the worst pandemic since the 1918 influenza. Thus, there is a critical need to create a vaccine that will protect against primary infection and re-infection with the virus (Amanat and Krammer, 2020).

The symptoms of SARS-CoV-2 infection include headache, fever, chills, and a persistent dry cough. Additional symptoms may include gastrointestinal distress, diarrhea, vomiting and loss of smell or taste. The main transmission route of SARS-CoV-2 is via respiratory exposure although infection may also occur via fecal-oral and ocular exposure. The causes of morbidity and mortality include pneumonia and the damaging cytokine storm elicited by infection, as well as multi-organ viral dissemination and organ failure in severe cases (Lippi et al., 2020). Close person-to-person contact via virus-containing droplets and aerosols is the primary mode of SARS-CoV-2 transmission. Importantly, since asymptomatic individuals also transmit the infection (Bai et al., 2020; Gandhi et al., 2020) there remains a need for vaccines that eliminate the viral load in the respiratory tract and prevent the development of a reservoir of asymptomatic individuals who serve as a silent source for transmission.

Emergency use authorization and subsequent FDA approval of mRNA vaccines, delivered intramuscularly, has altered the pandemic landscape, providing protection against severe disease and reducing mortality (Case et al., 2021). These vaccines generate strong systemic neutralizing antibody responses that limit viral infection. Some studies have detected mucosal IgA and IgG following immunization of non-human primates and humans with mRNA vaccines (Ketas et al., 2021; Mades et al., 2021; Tostanoski et al., 2021). However, it is unclear whether these vaccines elicit tissue-resident memory T cell responses (T_RM_) that are critical for preventing transmission and providing sustained protection against disease. Another important consideration in vaccine design is the possibility of vaccine associated enhanced respiratory disease (VAERD), which is the development of a more severe form of disease manifested after vaccination against the causative agent (DiPiazza et al., 2021; Halstead and Katzelnick, 2020; Liang et al., 2020; Seephetdee et al., 2021; Sekine et al., 2020) VAERD, which correlates with Th2 immune responses, is elicited by whole cell inactivated vaccines or Th1/Th2 skewing adjuvants such as alum. VAERD is reported as a side effect for inactivated vaccines against multiple pathogens, including respiratory syncytial virus (Graham et al., 1993; Kim et al., 1969), measles (Fulginiti et al., 1967; Nader et al., 1968), and SARS-CoV and MERS-CoV which are closely related to SARS-CoV-2 (Agrawal et al., 2016; Czub et al., 2005; See et al., 2006; Tseng et al., 2012). Thus, there remains a need for novel vaccines and immunization regimens that generate systemic and tissue resident immune responses, while limiting respiratory pathology.

Here we tested a subunit vaccine containing the soluble stabilized prefusion Spike (S) protein of SARS-CoV-2 adjuvanted with the Th1/2 polarizing adjuvant alum (aluminum hydroxide) alone or combined with the Th1/17 polarizing adjuvant Bordetella Colonization Factor A (BcfA) (Jennings-Gee et al., 2018). BcfA is an outer-membrane protein of the animal pathogen *Bordetella bronchiseptica* (Sukumar et al., 2009) that has adjuvant function, eliciting Th1/17 polarized systemic antibody and T cell responses to a variety of protein antigens (Jennings-Gee et al., 2018).

We leveraged the ability of alum to induce a strong and safe systemic immune response (Grifoni et al., 2020) with the ability of BcfA to attenuate alum-activated Th2 polarized immunity, to avoid enhanced respiratory disease (Bolles et al., 2011; Grifoni et al., 2020; Honda-Okubo et al., 2015; McPherson et al., 2016; S et al., 1993; Yasui et al., 2008). To generate both systemic and mucosal immunity, we employed a prime-pull immunization regimen in mice with intramuscular (i.m.) priming followed by an intranasal (i.n.) boost (Roces et al., 2019). This regimen generated S-specific IgG and IgA in the serum and respiratory tract, with a higher ratio of mucosal IgG2/IgG1 compared to the alum-adjuvanted vaccine, as we previously observed with BcfA-adjuvanted *Bordetella* vaccines (Jennings-Gee et al., 2018). Vaccines containing BcfA induced Th17 polarized CD4+ T_RM_ while, alum-adjuvanted vaccines, generated Th2 polarized systemic and mucosal CD4+ T cell responses. Importantly, immunized mice challenged with mouse-adapted SARS-CoV-2 were protected against both weight loss, and viral replication in the upper and lower respiratory tract. In addition, the BcfA-adjuvanted vaccine efficiently protected the respiratory tract against infection-associated lung damage, while the vaccine adjuvanted with alum alone did not. IL-17 knockout mice immunized with the same vaccine formulation and immunization regimen maintained high viral titer in the upper and lower respiratory tract and displayed respiratory pathology, suggesting that IL-17+ T cell responses were critical for protection by the BcfA-adjuvanted vaccine. Thus, the Th17 polarized mucosal and systemic T cell response, along with strong neutralizing antibodies generated by systemic priming with an alum-BcfA adjuvanted vaccine and mucosal booster with a BcfA-adjuvanted vaccine prevented SARS-CoV-2 induced severe illness, virus replication, and respiratory pathology.

## RESULTS

### Intramuscular priming with Spike/alum/BcfA (S/A/B) and intranasal boost with Spike/BcfA (S/B) elicits Th17 polarized mucosal and systemic T cell responses

We tested a subunit vaccine containing the soluble prefusion S protein with six stabilizing proline substitutions (Hsieh et al., 2020) as the antigen, adjuvanted with alum alone, or alum with BcfA.

We immunized C57BL/6 mice i.m. with 1 µg of S adsorbed to alum (S/A), or S protein with alum and 10 µg BcfA (S/A/B). We previously established that this combination dosage of adjuvants elicits protective immunity in a murine model of *B. pertussis* infection (Jennings-Gee et al., 2018). On d28, mice were boosted i.n. with S alone, S/A, or with S/B and evaluated 14 days post-boost (Figure 1A). To distinguish between tissue-resident and circulating T cells in the lungs, we injected mice intravenously (i.v.) with anti-CD45-PE 10 minutes prior to euthanasia (Anderson et al., 2014; Wilk et al., 2017) Lungs were enzymatically dissociated, single cell suspensions were stimulated with PMA/Ionomycin, then stained with antibodies against CD3, CD4 or CD8, CD44, CD62L, and CD69. Cells were fixed, permeabilized and stained with anti-IFNγ, anti-IL-17 and anti-IL-5 antibodies.

**Figure 1:**
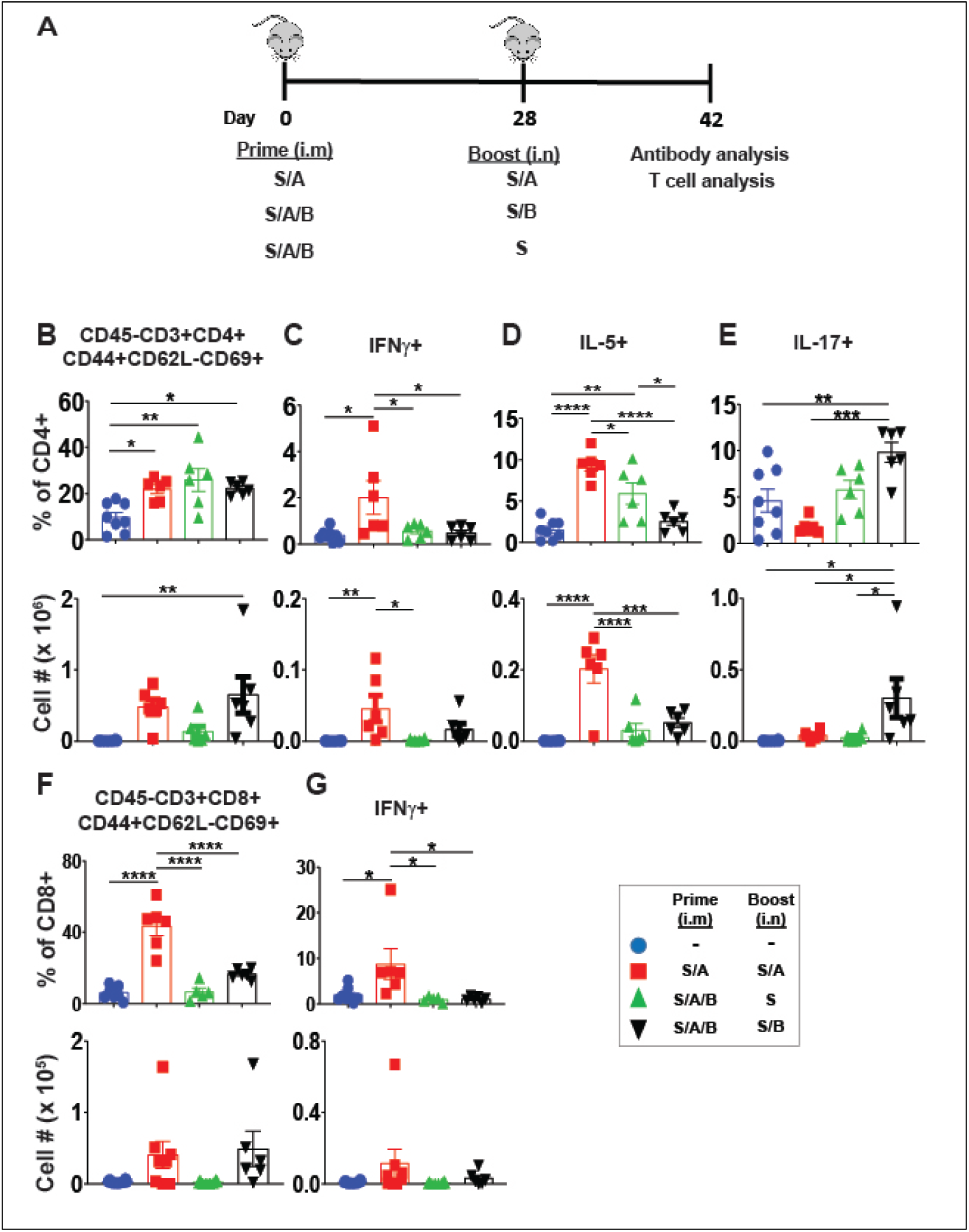
BcfA mucosal booster elicits Th17 polarized resident CD4+ lung memory T cells and attenuates Th2 responses elicited by alum. (A). Prime-pull immunization strategy to elicit mucosal immune response in C57/BL mice. Mice (6/group) were injected i.m. on d0 and inoculated i.n. on d28 with various vaccine formulations, with analysis of T cell populations in the lungs conducted on d42 (2 weeks post-boost). Mice were injected i.v. with CD45-PE antibody 10 min before euthanasia to distinguish between resident (CD45-) and circulating (CD45+) T cells. (B) The percentage and number of CD45-CD3+CD4+CD44+CD62L-CD69+ antigen-experienced CD4+ TRM, and (C) the percentage and number of IFNγ+ (D), IL-5+ (E) and IL-17+ cells is shown. Gating strategy in Figure S1. (F) The percentage and number of CD45-CD3+CD8+CD44+CD62L-CD69+ antigen-experienced CD8+ TRM and (G) IFNγ+ subset is shown. One-way ANOVA with Tukey’s multiple comparisons was used to detect differences between all experimental groups. Significance is indicated above for each group (∗p < 0.05; ∗∗p < 0.01; ∗∗∗p < 0.001; ∗∗∗∗p < 0.0001). The data are representative of 2 independent experiments. Mean and SEM are displayed.

Compared to naïve mice, the percentage of CD45-CD3+CD4+CD44+CD62L-CD69+ tissue-resident memory T cells (T_RM_) increased in all vaccinated groups (Figure 1B, see Figure S1 for the gating strategy). Although the total number of CD4+ T_RM_ in the lungs were higher than naïve mice in both S/A and S/A/B immunized groups, only the group primed with S/A/B and boosted with S/B showed a statistically significant increase (Figure 1B). Mice primed and boosted with S/A produced IFNγ (Figure 1C) and IL-5 (Figure 1D), while mice primed with S/A/B and boosted with S/B produced primarily IL-17 (Figure 1E). Notably, inclusion of BcfA in the vaccine significantly attenuated the proportion and number of IL-5 producing CD4+ T_RM_ (Figure 1D). The percentage and number of CD8+ T_RM_ (Figure 1F) that produced IFNγ (Figure 1G) increased in S/A immunized mice, but not in mice primed with S/A/B and boosted with S/B, suggesting that BcfA primarily elicited CD4+ T cells.

To understand the systemic T-cell response to vaccination we next examined circulating (CD45+) memory T cells from the same lung single cell suspensions. Mice immunized with either vaccine had a higher percentage of CD45+CD4+CD44+CD62L-CD69+ T cells in the lungs compared to naive mice while the number of these cells significantly increased in mice primed with S/A/B and boosted with S/B (Figure 2A). The T cells of mice primed and boosted with S/A produced IFNγ (Figure 2B) and IL-5 (Figure 2C), but negligible IL-17 (Figure 2D), indicating generation of a Th1/2 polarized immune response. In mice primed with S/A/B and boosted with S/B circulating memory CD4+ T cells did not produce IFNγ (Figure 2B) or IL-5 (Figure 2C) and only produced IL-17 (Figure 2D). Compared to naïve mice, the percentage and number of CD45+CD8+CD44+CD62L-CD69+ T cells increased with either alum or alum and BcfA-containing vaccines (Figure 2E). However, the proportion and number of IFNγ-producing cells only increased in S/A immunized mice (Figure 2F).

**Figure 2:**
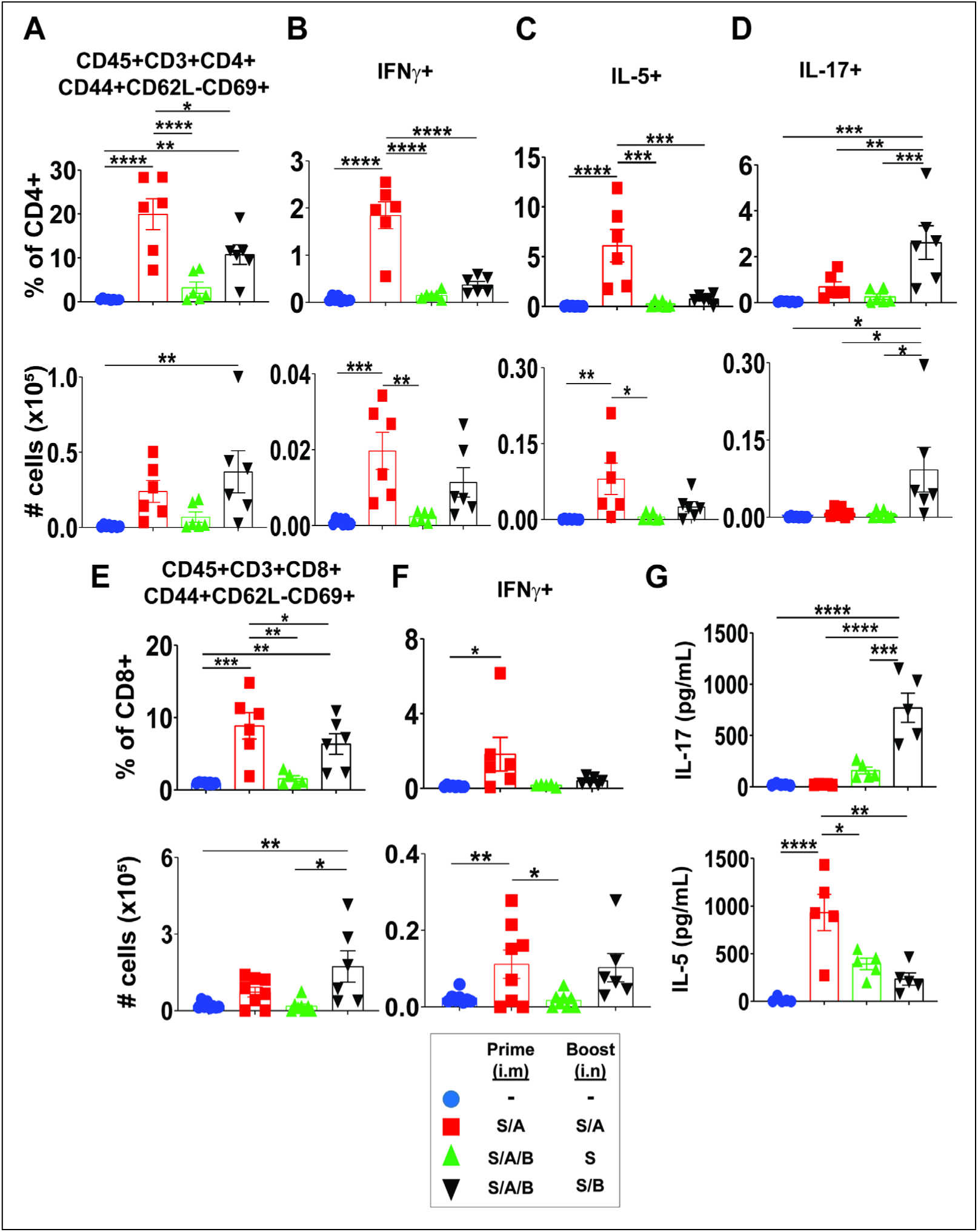
BcfA-adjvanted vaccines elicit Th17 polarized systemic T cell immunity. Lung cells (from mice in Figure 1 (6/group) were analyzed for the presence of CD45+ circulating T cells. (A) The percentage and number of CD45+CD3+CD4+CD44+CD62L-CD69+ antigen-experienced CD4+ T_RM_. (B) The percentage and number of IFNγ+ (C) IL-5+ and (D) IL-17+ cells is shown. (E) The percentage and of CD45+CD3+CD8+CD44+CD62L-CD69+ antigen-experienced CD8+ T_RM._ (F) The percentage and number of IFNγ+ subset is shown. (G) Splenocytes were isolated and processed into single cell suspensions before stimulation with 1μg S protein. IL-17 and IL-5 present in the culture supernatant 3 days post-stimulation was determined by ELISA. Differences between all experimental groups were analyzed by one-way ANOVA with Tukey’s multiple comparisons. Significance is indicated above for each group (∗p < 0.05; ∗∗p < 0.01; ∗∗∗p < 0.001; ∗∗∗∗p < 0.0001). Mean and SEM are shown.

We then tested the cytokines produced by spleen cells of immunized mice, following stimulation with S protein for 72 hr. Spleen cells of mice primed with S/A/B and boosted with S/B produced IL-17 (Figure 2G), but negligible IL-5 (Figure 2H). Spleen cells of S/A primed and boosted mice primarily produced IL-5 (Figure 2H) while IFNγ was not detected in supernatants of any of the groups (data not shown). Thus, S/A/B prime and S/B boost elicited Th17 polarized systemic responses, over-riding the Th2 skewed responses elicited by alum.

We then evaluated whether immunization with BcfA-containing vaccines generated antigen-specific T_RM_ cells. Dissociated lung cells were stimulated with peptide pools derived from S (Miltenyi Biotec) for 6 hr and stained as described above. The percentage and number of antigen-experienced cells increased in mice primed with S/A/B and boosted with S/B (S/A/B_S/B) (Figure 3A, gating strategy in Figure S2). These cells did not produce IFNγ (Figure 3B) or IL-5 (Figure 3C) and produced only IL-17 (Figure 3D). The proportion and number of antigen-experienced and cytokine-producing cells did not increase in mice immunized with S/A/B and boosted with S alone (S/A/B_S), compared to naïve mice or mice that received the S/B booster. Thus, the inclusion of BcfA in the mucosal boost generated antigen specific T_RM_ in the lungs. Interestingly, while the proportion and number of antigen-specific circulating (CD45+) memory T cells increased (Figure 3E), changes in percentage and number of cytokine-producing cells did not reach statistical significance. This result suggests that the antigen-specific cells were largely tissue-resident. Furthermore, this vaccine did not elicit antigen specific CD8+ T cells (data not shown).

**Figure 3:**
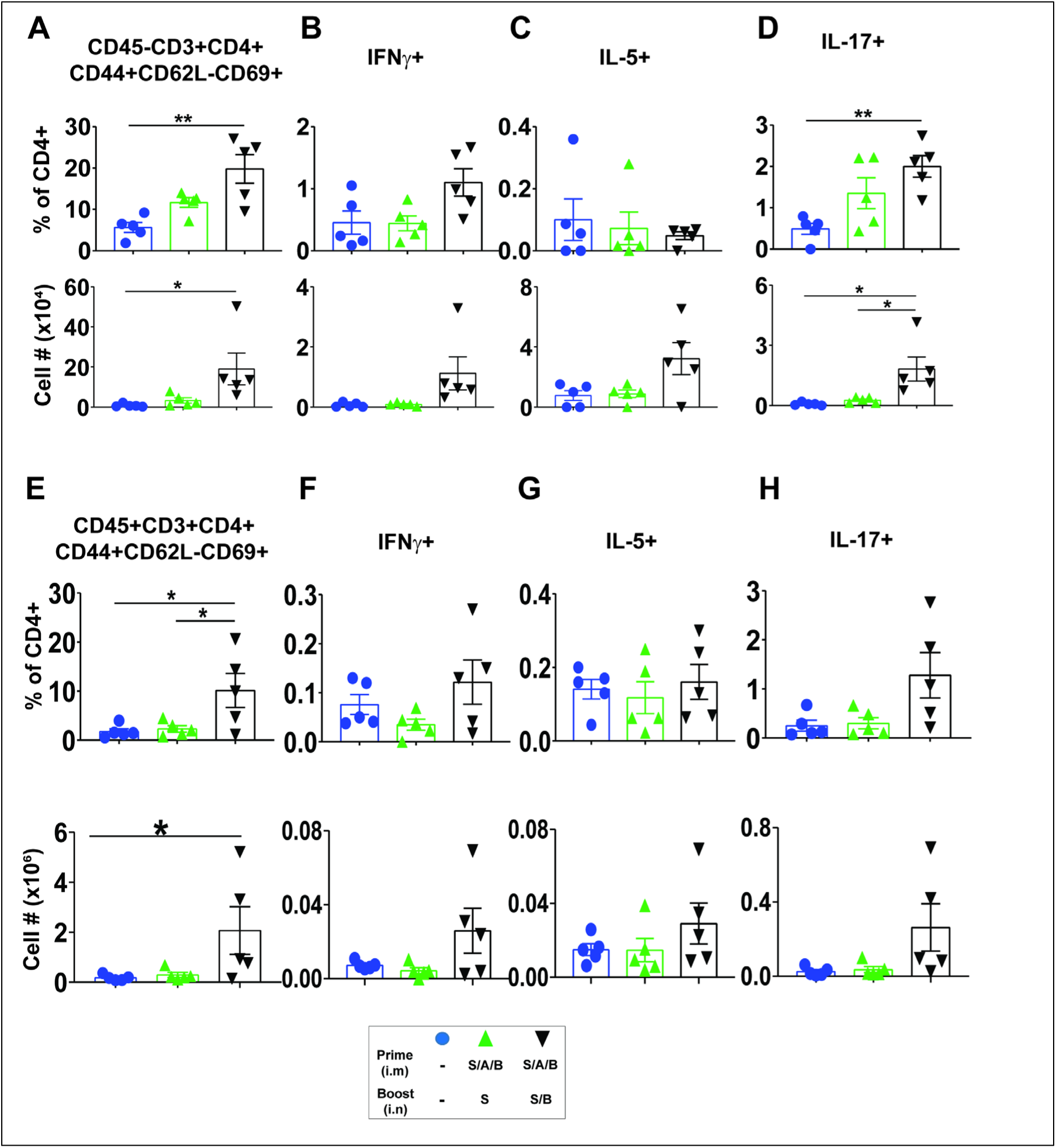
BcfA-adjuvanted mucosal booster elicits S-specific Th17 polarized lung resident CD4+ T_RM_ memory T cells. Lung cells were stimulated for 6 hours with S-derived peptide pools (S1 and S2) at a concentration of 1μg/ml in the presence of protein transport inhibitors. Surface staining and ICS was conducted to analyze both CD45-resident and CD45+ circulating T cells. (A) The percentage and number of CD45-CD3+CD4+CD44+CD62L-CD69+ antigen-experienced CD4+ T_RM_. (B) The percentage and number of IFNγ+ (C) IL-5+ and (D) IL-17+ cells is shown. (E) The percentage and of CD45+CD3+CD4+CD44+CD62L-CD69+ antigen-experienced CD4+ T_RM._ (F) The percentage and number of IFNγ+ (G) IL-5+ and (H) IL-17+ subsets is shown.. One-way ANOVA with Tukey’s multiple comparisons was used to detect differences between all experimental groups. Significance is indicated above for each group (∗p < 0.05; ∗∗p < 0.01; ∗∗∗p < 0.001; ∗∗∗∗p < 0.0001). N= 5 animals per group, mean and SEM of the results are displayed.

Thus, the addition of BcfA to the priming immunization and a booster with BcfA as the sole adjuvant elicited Th17 polarized systemic and mucosal CD4+ T cells and significantly attenuated IL-5 responses primed by alum.

### Prime-pull immunization generated Tfh and GC B cells in the draining lymph nodes

T follicular helper (Tfh) cells are a specialized helper T cell subset that play a critical role in support of antibody production by B cells (Crotty, 2014). We determined whether our prime-boost immunization regimen and vaccines generated Tfh cell populations. Indeed, we observed Tfh (PD-1^hi^CXCR-5^hi^) cells in the draining mediastinal lymph nodes of mice immunized with S/A alone or primed with S/A/B and boosted with S/B (Figure 4A). As Tfh populations support B cell activation and neutralizing antibody production, we also determined whether our immunization regimen and vaccines supported the activation of germinal center (GC) B cells (Cyster and Allen, 2019; Krautler et al., 2017). All adjuvanted vaccines generated activated (Fas+GL-7+) GC B cells (Figure 4B) compared to naïve mice, as also observed in mice immunized i.m. only with an alum-adjuvanted COVID-19 vaccine (DiPiazza et al., 2021). Figure S3 shows the gating strategy. Together, these data suggest that systemic priming with an alum-containing vaccine activates Tfh and GC B cell responses that promote S-specific antibody production.

**Figure 4:**
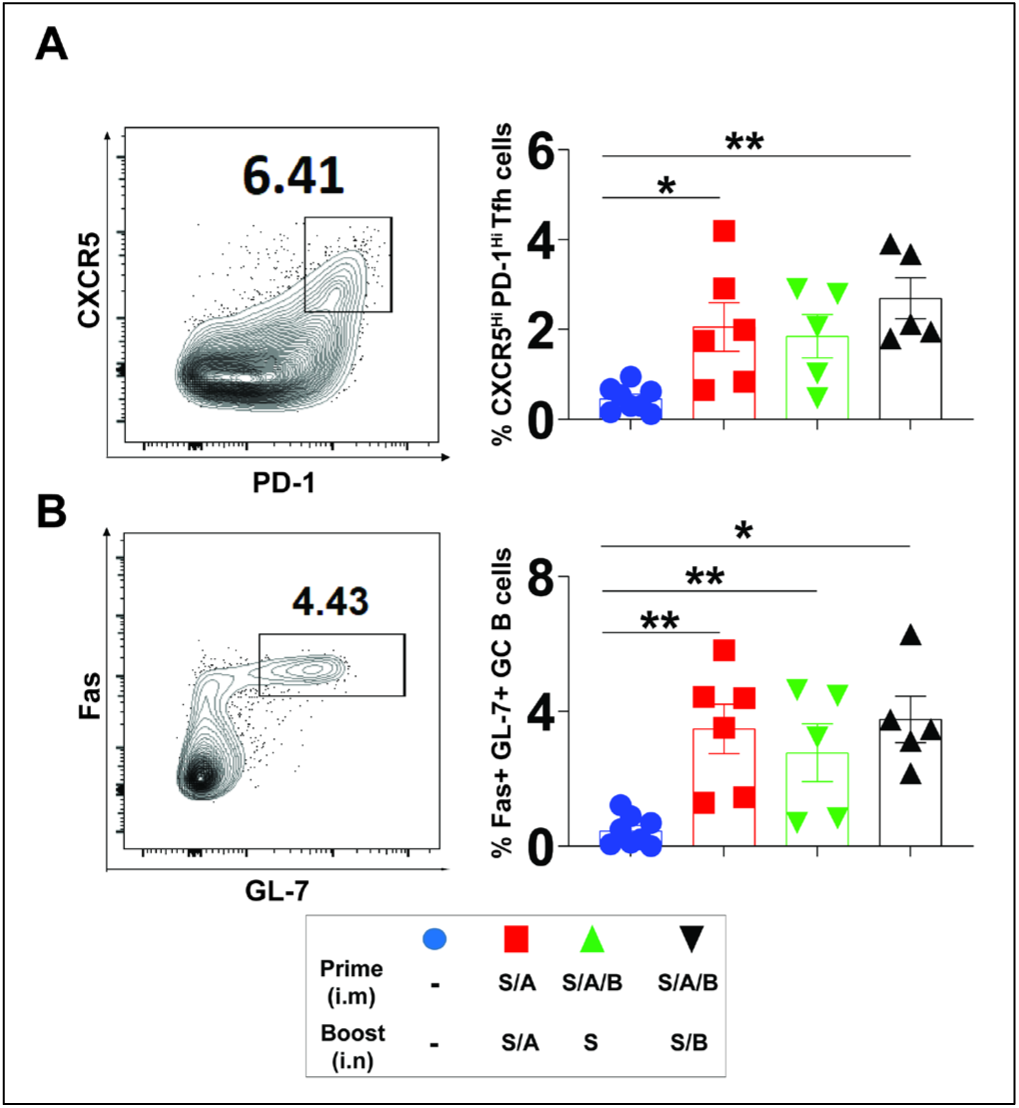
Immunization with alum or alum+BcfA adjuvanted vaccines generates PD-1^high^ CXCR5^high^ (Tfh) cells and Fas+GL7+ (GC B) cells in the draining mediastinal lymph node. Single cell suspensions of the draining (mediastinal) lymph nodes were evaluated for presence of T_FH_ and germinal center B cell populations. T_FH_ populations were identified as activated (CD44+) PD-1^hi^CXCR5^hi^ cells. Germinal center B cell populations were identified as Fas+GL-7+ cells. (A) Percentage of T_FH_ out of total CD4+ cells. (B) Percentage of GC B cells out of total CD19+ B cells. One-way ANOVA with Tukey’s multiple comparisons was used to detect differences between all experimental groups. ∗p < 0.05; ∗∗p < 0.01; ∗∗∗p < 0.001; ∗∗∗∗p < 0.0001. N=6 animals per group. Data are displayed as mean and SEM and are representative of 2 independent experiments.

### BcfA-adjuvanted vaccines elicit Th1 polarized mucosal and systemic IgG antibodies

Generation of S-specific antibodies by vaccination is correlated with protection against SARS-CoV-2 (DiPiazza et al., 2021; Seephetdee et al., 2021). We determined whether the prime-pull immunization regimen generated both mucosal and systemic antibodies. We quantified the S-specific IgG and IgA antibodies in the lungs of naive and immunized mice. All immunized groups of mice had higher IgG antibodies in the lungs compared to naive mice (Figure 5A). S/A and S/B boosted mice had higher antibodies than mice boosted with S alone, suggesting that an adjuvant is important for generating mucosal antibodies. Additionally, the avidity of S-specific IgG, determined by addition of sodium thiocyanate (Macdonald et al., 1988), was higher in the mice that received the S/B booster compared to booster with S alone (Figure 5B). We then tested whether the addition of BcfA altered the Th1/Th2 polarization of the mucosal antibodies. Mice boosted with S/B had a higher ratio of IgG2b/IgG1 compared to booster with S alone and higher IgG2c/IgG1 compared to mice primed and boosted with S/A or primed with S/A/B and boosted with S alone (Figure 5C). Thus, the combination of alum and BcfA as adjuvants increased mucosal IgG antibody avidity and skewed the overall IgG response towards the Th1 phenotype.

**Figure 5:**
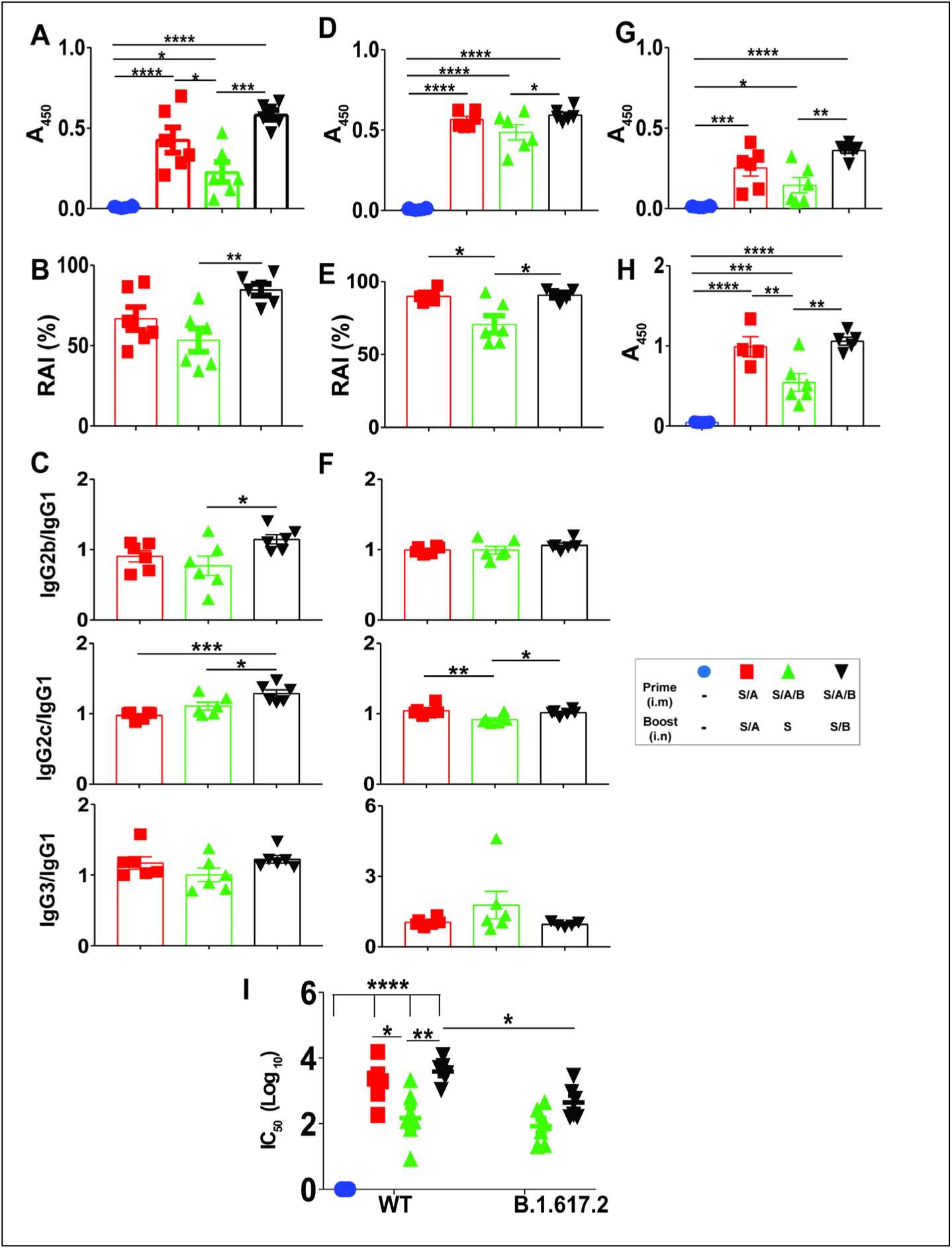
BcfA mucosal booster generates Th1-polarized mucosal and systemic antibodies. ELISA analysis evaluated S-specific antibodies in lung homogenates and serum. (A) S-specific IgG in lung homogenates at 1:1250 dilution. (B) 2M sodium thiocyanate (NaSCN) was added to the ELISA to measure antibody avidity of S specific IgG in lung homogenates. Relative avidity index (RAI) was calculated: (A450 at 2M) / (A450 at 0 M) *100. (C) S-specific IgG isotypes in lung homogenates were determined using specific secondary antibodies. The ratio of each IgG2 subclass to IgG1 is shown. (D) S-specific IgG in serum at 1:12500 dilution. (E) Relative avidity index of serum IgG was calculated with the addition of 2M sodium thiocyanate in the ELISA. (F) The ratio of each IgG2 subclass to IgG1 in serum. (G) IgA in lung homogenates at 1:50 dilution. (H) IgA in serum at 1:12500 dilution. (I) A Gluc-based lentiviral SARS-CoV-2 pseudovirus expressing the WT S protein or the B.1.617.2 (Delta) variant was used to determine the neutralizing activity of serum antibodies. Differences between experimental groups were determined by one-way ANOVA with Tukey’s multiple comparisons. Significance is indicated above for each group (∗p < 0.05; ∗∗p < 0.01; ∗∗∗p < 0.001; ∗∗∗∗p < 0.0001). N= 6 animals per group. Data are representative of two independent experiments. Mean and SEM of the results are displayed.

We then quantified the relative amounts of S-specific IgG in the serum. All groups immunized with adjuvanted vaccines had specific serum antibodies at levels higher than naïve mice (Figure 5D). The relative amount of antibody was higher in mice that received either S/A or S/B in the booster compared to S alone (Figure 5D), suggesting that an adjuvant in the mucosal booster increased the amount of circulating IgG antibodies and IgG antibody avidity (Figure 5E).

### Prime-pull immunization generated mucosal and systemic IgA

We found that immunization with S/A or S/A/B_S/B generated IgA in the lungs (Figure 5G) and in the serum (Figure 5H), demonstrating that the mucosal booster induces IgA locally and systemically. In contrast, systemic immunization with alum-adjuvanted S containing vaccines does not generate circulating IgA responses (DiPiazza et al., 2021). Thus, IgA elicited by a mucosal booster also enters the circulation.

### Serum antibodies elicited by S/A/B vaccine neutralized the wildtype SARS-CoV-2 and Delta variant

We tested the ability of serum antibodies to neutralize a pseudotyped lentivirus bearing S protein of SARS-CoV-2 WA1 strain. Mice immunized with S/A, or BcfA-containing vaccines had high neutralizing activity (Figure 5I) compared to mice boosted with S alone. Antibodies elicited by BcfA-adjuvanted vaccines also neutralized the virus pseudotyped with S from the SARS-CoV-2 lineage B.1.167.2 (Delta variant), suggesting that the BcfA-adjuvanted vaccine elicits a broad polyclonal antibody response with protection against an early isolate of SARS-CoV-2 as well as a currently circulating VOC.

### Immunization of mice with S/A or S/A/B prevented SARS-CoV-2 infection associated weight loss and reduced viral titers in the respiratory tract

To determine whether our immunization regimens prevented virus infection *in vivo*, we challenged naive and immunized mice i.n. with 5×10^4^ PFU of mouse-adapted SARS-CoV-2 (strain MA10) (Figure 6A). We recorded daily body weight in one cohort of infected mice until d10 p.i. to assess whether immunization affected disease severity. Naive mice and mice immunized with S alone lost 10-15% body weight beginning on d2 post-infection and recovered to pre-challenge weight by d8 p.i. (Figure 6B). In contrast, mice immunized and boosted with S/A, or immunized with S/A/B and boosted with S/B did not lose weight (Figure 6B), indicating that the vaccines prevented severe disease. We quantified viral load in the nose and lungs on d2 p.i. in a second cohort infected at the same time. Viral load in the nose (Figure 6D) and lungs (Figure 6E) was 5-6 logs lower in both vaccinated groups, compared to S-immunized or naïve mice. Thus, a vaccine adjuvanted with alum alone, or alum and BcfA, prevented virus-associated weight loss and virus infection of the upper and lower respiratory tract. To test the contribution of Th17 polarized T cells to this response, we immunized IL-17 knockout mice i.m. with S/A/B and boosted the mice i.n. with S/B 28 days later. Naïve and immunized knockout mice were challenged with MA10 virus 14 days later. S/A/B_S/B immunized IL-17 KO mice did not lose weight, suggesting that protection against severe disease was unaffected by the absence of IL-17+ T cells. However, viral load in the nose of IL-17 KO mice was similar in naïve and immunized mice (Figure 6F) and only reduced by ∼ 1 log in the lungs of immunized mice (Figure 6G). This may be attributed to the lower viral load in the lungs of IL-17 KO mice compared to wildtype mice. These data suggest that the Th17 polarized T cell response generated by the BcfA-adjuvanted vaccine is critical for clearing the virus from the respiratory tract.

**Figure 6:**
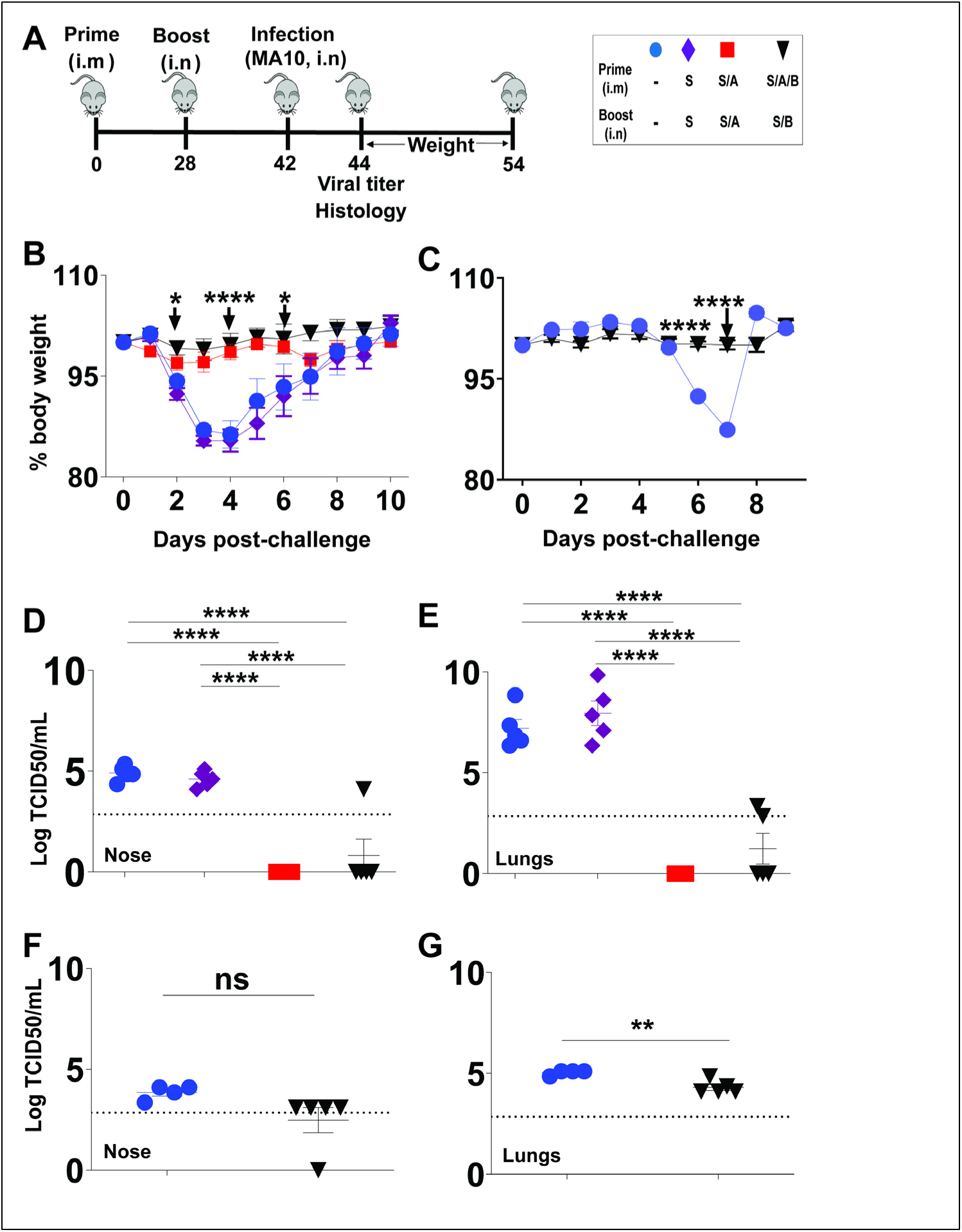
Prime-pull immunization with either alum or BcfA adjuvanted vaccines protects from SARS CoV2 MA10 induced disease and reduces viral infection of the nose and lungs. (A) Prime-pull immunization strategy to elicit mucosal immune response in C57/BL or IL-17 KO mice. At 2 weeks post-boost mice (10 /group) were challenged with the mouse adapted SARS CoV2 strain (MA10). The percent of original body weight on each day post-challenge was calculated through day 10 post infection (5 mice/group) in (B) C57BL/6 mice and (C) IL-17 KO mice. Two-way ANOVA with Tukey’s multiple comparisons was used to detect differences between all experimental groups in B and Student’s t-test in C. Viral titer expressed as log Median Tissue Culture Infectious Dose (TCID50) was quantified from nasal septum (D) and lung homogenates (E) of C57BL/6 mice and nasal septum (F) and lungs (G) of IL-17 KO mice obtained on day 2 post infection (4-5 mice/group). One-way ANOVA with Tukey’s multiple comparisons was used to detect differences between experimental groups in D and E, and Student’s t-test in F and G. Mean and SEM of the results are displayed. Significance is indicated above for each group (∗p < 0.05; ∗∗p < 0.01; ∗∗∗p < 0.001; ∗∗∗∗p < 0.0001).

### Marked lymphocyte and PMN infiltrate without epithelial damage following SARS-CoV-2 challenge in the lungs of mice primed with S/A/B and boosted with S/B

We evaluated formalin-fixed paraffin embedded lung sections harvested on d2 p.i. to determine the extent of pneumonia, immune cell infiltration and epithelial damage. Histopathological evaluation of sections stained with hematoxylin and eosin detected marked hemorrhage (filled arrows) and edema (open arrows) around blood vessels and airways in naïve challenged mice (Figure 7A) and mice immunized with S alone (Figure 7B). Mixed eosinophilic proteinaceous debris was often admixed, sometimes near vessels rimmed by hemorrhage. Significant type II pneumocyte hyperplasia (asterisks) was evident in affected areas, resulting in moderate to marked thickening of alveolar walls, often accompanied by infiltration of large macrophages. Bronchioles exhibited degeneration and necrosis (ovals), with necrotic cellular debris, and blebbing apical epithelial cell debris. Mice primed and boosted with S/A displayed a focal pattern of hemorrhage and edema with milder degeneration and airway necrosis (Figure 7C). Large foamy macrophage infiltration was observed with some lymphocyte aggregates (tailed arrows) in perivascular spaces with little edema. Eosinophilic debris was less evident in this group. Mice primed with S/A/B and boosted with S/B (Figure 7D) showed comparably mild thickening of alveolar walls, mild degeneration of bronchiolar epithelium, some perivascular and vascular inflammation, and marked increase in perivascular and peribronchiolar lymphocytic aggregates with mitoses (tailed arrows), suggesting expansion of the lymphoid population. The absence of epithelial damage was notable in this group. Semi-quantitative scored showed reduced thickening of alveolar walls (Figure 7E) reduced alveolar macrophages (Figure 7F) and reduced degeneration and necrosis (Figure 7G) in mice immunized with S/A/B_S/B compared to naïve challenged mice. Lymphocytes were increased in this group compared to naïve challenged mice (Figure 7H). Total cellularity, edema and hemorrhage were similar between naïve and immunized mice (Figure S4). We then tested the presence of SARS-CoV-2 N protein as a second measure of virus presence in the lungs. IHC analysis revealed strong and widespread N protein staining in epithelial cells of naïve challenged mice (Figure 7I) and mice immunized with S alone (Figure 7J). Mice primed and boosted with S/A showed foci of N protein staining in 2 of 5 mice (Figure 7K and Table S1), suggesting that protection from infection was inefficient in this group. Remarkably, N protein staining was completely absent from the lungs of all mice immunized with S/A/B and boosted with S/B (Figure 7L and Table S1), suggesting either that this vaccine combination prevented infection more efficiently, or that the more robust immune cell infiltrate accelerated clearance of virally infected cells and debris. Thus, histopathological and IHC analysis revealed that priming mice with S/A/B and boosting with S/B prevented viral infection and respiratory disease. Histopathology of lungs from naïve and immunized IL-17 KO mice showed milder pathology overall compared with wild-type mice. Naïve challenged mice displayed thickening of the alveolar wall with hemorrhage, type II pneumocyte hyperplasia and inflammation, degeneration and sloughing in the airways with an inflammatory infiltrate and tissue necrosis (Figure S5A). Lungs of immunized IL-17KO mice also displayed hemorrhage, necrosis, airway degeneration, and type II pneumocyte hyperplasia. Lymphoid expansion, congestion and hypercellularity was also observed (Figure S5B). Histopathology quantification showed a slight increase in alveolar wall thickening (Figure S5C) and no change in alveolar macrophages (Figure S5D) or necrosis (Figure S5E) in immunized mice. Infiltration of lymphocytes and plasma cells was significantly higher (Figure S5F) in immunized mice compared to naïve mice, suggesting that while the immunization recruited lymphocytes to the tissues, these did not have antiviral function. These data show that while IL-17 KO mice had milder respiratory disease compared to wild-type C57BL/6 mice, immunized mice were not protected from respiratory pathology. Thus, Th17 polarized T cells generated by BcfA-adjuvanted vaccines contribute to prevention of respiratory damage following SARS CoV-2 challenge. IHC did not detect N protein staining in lungs of IL-17 KO naïve or immunized mice (data not shown), likely due to the overall lower viral titer in the respiratory tract compared with wild-type C57BL/6 mice.

**Figure 7:**
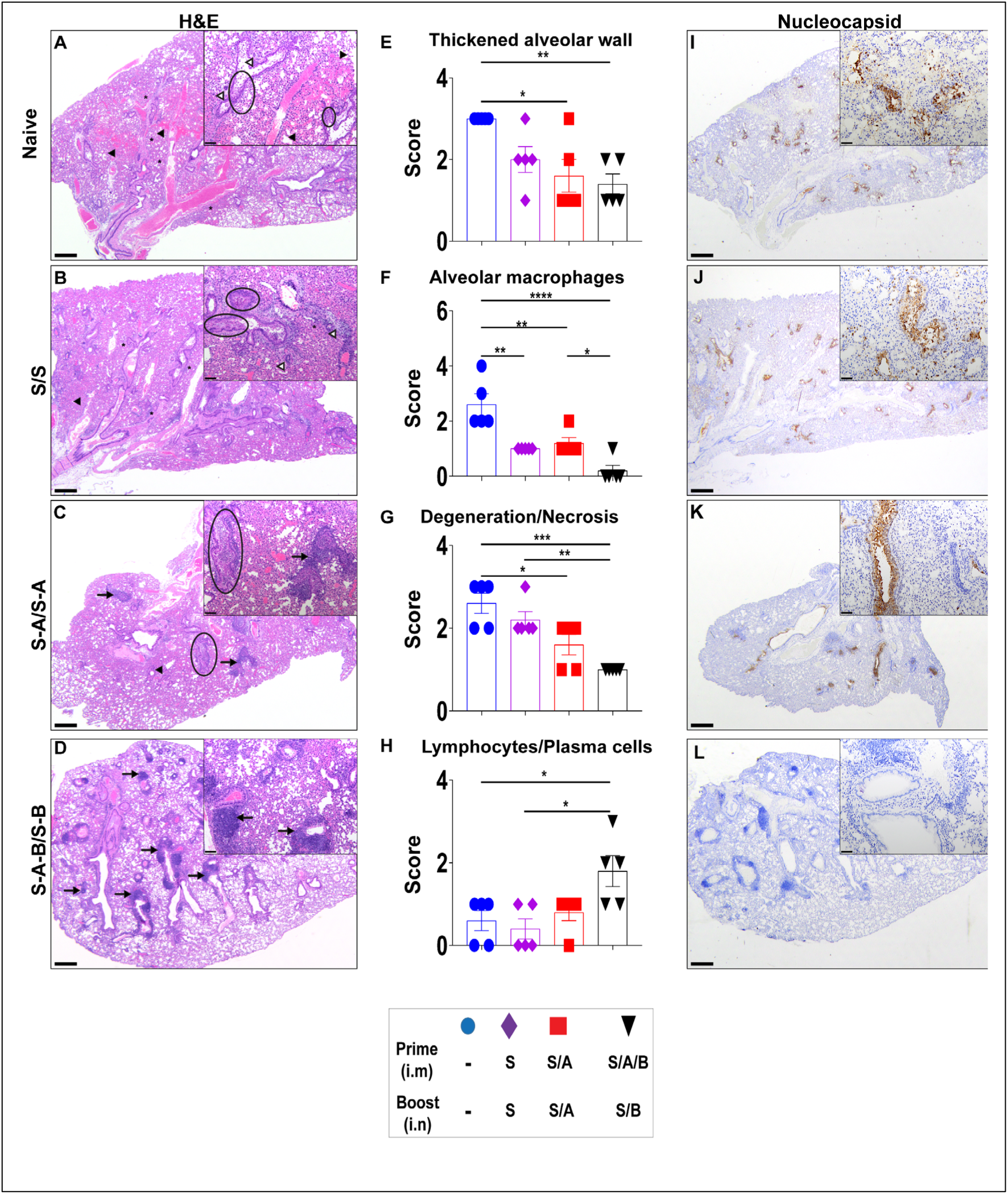
Mice immunized with alum and BcfA adjuvanted vaccines have reduced lung pathology and viral antigen compared to alum adjuvanted vaccines after challenge with SARS-CoV-2 MA10. Formalin fixed, paraffin embedded lungs harvested on d2 post-infection were sectioned and stained with H&E (A-D) to evaluate inflammation and tissue damage. (E-H). Semi-quantitative scoring of (E) alveolar wall thickness (F) presence of alveolar macrophages (G) degeneration and necrosis (H) presence of lymphocytes and plasma cells. (I-L) Nucleocapsid viral antigen was evaluated by IHC. H&E staining: 2x magnification (scale bar 500 µm) with 10x inset (scale bar 100 µm); Nucleocapsid staining 2x magnification with 20x inset (scale bar 50 µm).

## DISCUSSION

In this study, we identified an immunization strategy and vaccine formulation that elicits protective immunity against SARS-CoV-2 both systemically and at the infection site. Generation of systemic immunity is important for preventing viral infection and dissemination while generation of mucosal immunity is critical for clearance of pathogens and infected cells from the respiratory tract (Matuchansky, 2021; Russell et al., 2020).

The S protein is a target during natural SARS-CoV-2 infection, eliciting S-specific antibodies and T cells (Grifoni et al., 2020; Hellerstein, 2020; Le Bert et al., 2020; Peng et al., 2020; Tan et al., 2020; Wajnberg et al., 2020; Zeng et al., 2020). We used a highly stabilized form of the pre-fusion S protein as the antigen since it is produced more efficiently than the native S and it maintains its structure better than the original 2Pro stabilized form. As such, it elicits neutralizing antibodies with greater potency and broader specificity than post-fusion S and is the preferred antigen for new SARS-CoV-2 vaccines (Arunachalam et al., 2021; Hsieh et al., 2020).

We leveraged the noted strengths of alum in generating strong and safe systemic responses with the ability of BcfA to attenuate Th2 responses when combined with alum. I.m. priming and i.n. boosting with S/A generated systemic and mucosal T cells and antibodies. Notably, in contrast to alum-adjuvanted vaccines delivered i.m. only (Arunachalam et al., 2021; DiPiazza et al., 2021), prime-pull immunization with S/A produced IgA in the serum and lungs, demonstrating that changing the delivery route alters the composition of the immune response. Although alum and alum/BcfA-adjuvanted vaccines induced mucosal IgA and IgG, the IgG avidity and ratio of IgG2/IgG1 was higher in the lungs of mice primed with S/A/B, and boosted with S/B. In contrast, serum antibodies in both vaccine groups were similar in avidity and IgG2/IgG1 ratio, suggesting that the mucosal booster with S/B resulted in antibody maturation *in situ*.

CD8+ and CD4+ T cell memory responses specific for S as well as for structural proteins M and N are detected in COVID-19 convalescent individuals (Grifoni et al., 2020; Kared et al., 2021; Sekine et al., 2020), suggesting that cell-based immunity is critical for protection against SARS-CoV-2. S/A immunization, with alum as the sole adjuvant, generated CD4+IL-5+ circulating and T_RM_ cells, confirming the Th2 bias of alum-primed immunity. As we reported previously in murine models of *Bordetella pertussis* infection (Jennings-Gee et al., 2018), i.m. priming with alum/BcfA adjuvanted vaccines attenuated the systemic Th2 polarized responses primed by alum. Here we showed that mucosal booster with either S/A, or S/B elicited CD4+ T_RM_ T cells, correlated with long-lived protection against respiratory pathogens. Addition of BcfA induced IL-17-producing T cells with significant attenuation of IL-5. Circulating and splenic T cell responses mirrored the polarization displayed by T_RM_. Both alum and BcfA elicit CD8+ T cell responses against other antigens (Jennings-Gee et al., 2018). However, we detected minimal CD8+ T cell responses to the S protein, suggesting that T cell responses induced by the same adjuvant vary with the antigenic composition of the vaccine. These data further suggest that CD8+ T cells are not required for clearance of SARS-CoV-2 from the respiratory tract.

Post-mortem studies suggest that COVID-19 infection blunts Tfh responses (Kaneko et al., 2020) that are critical for induction of sustained humoral immunity. Prime-pull immunization with either alum or alum/BcfA adjuvanted vaccines generated Tfh cells. Th1 polarization of antibodies elicited by BcfA-adjuvanted vaccines implicate Th1 polarized Tfh cells in this response. Ongoing studies in our group are dissecting Tfh cell phenotypes and the longevity of the corresponding antibodies generated by alum or alum/BcfA-adjuvanted vaccines.

Systemic immunization is the approved route for most vaccines, including those against SARS-CoV-2, and generates strong serum antibodies and circulating T cell responses. However, subunit vaccines delivered systemically do not generate mucosal immunity. I.n immunization generates tissue-resident memory T cell responses that provide long-lived protection in the upper respiratory tract (Allen et al., 2018; Wilk and Mills, 2018). We reasoned that i.m. priming and i.n. booster would elicit strong systemic and mucosal responses and may be especially useful for booster vaccination of individuals already immunized by approved COVID-19 vaccines (Lavelle and Ward, 2021; Lycke, 2012; Russell et al., 2020). A similar strategy was tested in macaques that were primed i.m. with S+alum and boosted i.n. with a nanoparticle formulation containing S and CpG, poly I:C and IL-15 as adjuvants (Sui et al., 2021). This regimen reduced viral load in the upper and lower respiratory tract and, also supports the possibility that the combination of systemic and mucosal immunization will be the most effective in preventing COVID-19 infection.

Weight loss is a key manifestation of disease in mice challenged with SARS-CoV-2 (DiPiazza et al., 2021; Tostanoski et al., 2021). Mice immunized with either S/A or S/A/B_S/B did not lose a significant amount of weight after challenge, while naive mice did, only recovering normal body weight by day 8 post-challenge. Mice immunized with alum or alum/BcfA containing vaccines had low viral titer in the nose and lungs, suggesting that both formulations provide similar protection against severe disease. Although IL-17 KO mice immunized with S/A/B_S/B did not lose weight following virus challenge, viral load in the nose did not decline and declined slightly in the lungs, suggesting that IL-17-producing T cells are critical for the anti-viral response. Histopathology and IHC analysis revealed an important distinction. As reported by other studies (DiPiazza et al., 2021), S/A immunized mice had evidence of pneumonitis and epithelial damage and prolonged expression of nucleoprotein antigen, despite the production of mucosal IgA. In contrast, the lungs of mice immunized with BcfA-adjuvanted vaccines had a leukocyte infiltrate, little evidence of damage to the lung tissue and complete absence of nucleoprotein expression. Together our data show that a subunit vaccine that elicits Th1/17 polarized systemic and mucosal immunity is highly effective against SARS-CoV-2. In naïve IL-17 KO mice, although lung pathology was milder than in wild-type mice, immunized mice also displayed respiratory disease. Thus, the IL-17+ T cells produced by S/A/B_S/B contribute to reduction of viral load and protection against respiratory disease.

Multiple studies suggested a significant role for IL-17 in SARS-CoV-2 infection-induced immunopathology (Parackova et al., 2021; Wu et al., 2020; Xu et al., 2020) and argued for targeting IL-17 as a plausible therapeutic strategy (Orlov et al., 2020; Pacha et al., 2020; Sarmiento-Monroy et al., 2021; Wu et al., 2020). In addition, some proposed Th17 polarized responses to be the key player in VAERD instead of Th2, although current evidence is inconclusive (Hotez et al., 2020; Liang et al., 2020). Our results suggest that activation of Th17 responses while attenuating Th2 responses limits pathology and protects against COVID-19 infection. Testing of BcfA-adjuvanted vaccines in larger animal models to validate these results in mice will be necessary prior to clinical trials.

Together, our data show that prime-pull immunization with a combination of alum and BcfA as adjuvants, generates S-specific systemic and mucosal Th1/17 polarized immune responses that are highly effective at preventing SARS-CoV-2 infection, and preventing respiratory damage. Given that global vaccination rates are on the rise, there will be an increasing need for booster vaccines that extend protection provided by currently approved mRNA vaccines. Thus, it will be important to test whether i.n. booster with S/B generates mucosal immunity in individuals previously immunized with mRNA vaccines, and thereby increases the longevity of protection.

## MATERIALS AND METHODS

### Biosafety

All experiments were performed in accordance with standard operating procedures at Biosafety Level 2 (BSL-2) or Biosafety Level 3 (BSL-3) as appropriate. Work with live SARS-CoV-2 was performed in BSL-3 facilities according to standard operating procedures approved by the Ohio State University BSL-3 Operations Group and Institutional Biosafety Committee. Samples were removed from the BSL3 facility for IHC analysis after fixation and decontamination with 4% paraformaldehyde for a minimum of 1 h according to an in-house validated and approved method of sample decontamination.

### Mice

All experiments were reviewed and approved by The Ohio State University (OSU) Institutional Animal Care and Use Committee (protocol numbers 2020A00000054 and 2020A00000001). C57Bl/6J mice and IL-17 KO mice (male and female, older than 6 weeks old) were bred in-house.

### Reagents

BcfA was produced and purified as described previously (24) with stringent endotoxin removal. Endotoxin level was <5 EU/mg protein (51). Stabilized S protein containing six proline substitutions was produced and purified as described (Hsieh et al., 2020). Aluminum hydroxide (alum) and fetal bovine serum (FBS) was from Sigma-Aldrich (St. Louis, MO). ELISA kits were from eBioscience (Thermo Fisher Scientific). Flow cytometry antibodies were from eBioscience, BD Biosciences or R&D Systems (see Table 1).

### Isolation of mouse-adapted SARS-CoV-2 with an intact furin cleavage site

Mouse-adapted SARS-CoV-2 (strain MA10 generated by the laboratory of Dr. Ralph Baric, UNC) was obtained from BEI Resources,(Leist et al., 2020). and. The provided stock was found to contain a mixture of wild-type (intact furin cleavage site, RRAR) and a large proportion of viruses with mutations/deletions in the furin cleavage site. To isolate a pure stock of SARS-CoV-2 MA10 containing the intact furin cleavage site, virus was passaged once in primary, well differentiated human bronchial epithelial cells to enrich for viruses that could infect these primary cells. Progeny virus was harvested at day 5 post-infection, serially diluted 10-fold and used to inoculate Vero-TMPRSS2 cells in a plaque assay. Two days post inoculation, 50 individual small plaques were picked and used to inoculate one well of Vero-TMPRSS2 cells in a 12-well plate. The progeny virus from each well was harvested at day 2 post-inoculation and viral RNA was extracted from the harvested supernatant for RT-PCR with primers that amplify the region of the S gene encoding the furin cleavage site (forward: 5’-AAATCTATCAGGCCGGTAGC-3’; reverse: 5’-GAAGCCAGC ATCTGCAAGTG-3’). The PCR products were Sanger sequenced commercially using the forward primer. Four of 50 plaques were found to contain SARS-CoV-2 MA10 with the intact furin cleavage site. These viruses were grown in Vero-TMPRSS2 cells for another passage and sequenced again to confirm the presence of the intact furin cleavage site before being used in experiments.

### Immunizations

Mice were lightly anesthetized with 2.5% isoflurane/O_2_ for immunization. I.m. immunization on day 0 was delivered in 100 µL volume divided between both forelimbs. I.n. booster was inoculated in 50 µL divided between both nares on days 28-35. Acellular vaccines contained 1 µg stabilized S alone (S) or mixed with 10 µg BcfA (S/B). S protein was adsorbed to 130 µg of alum by rotating the suspension for 30 minutes at room temperature (S/A). Beads were pelleted and resuspended in 100 µL fresh PBS for i.m. injection or 50 µL for i.n. instillation. S/A/B included 10 µg BcfA and 1 µg S adsorbed to alum as above.

### SARS-CoV-2 S pseudotyped lentivirus neutralization assay

Virus neutralization by serum and lung antibodies was performed as described (Zeng et al., 2020). We used 100μL *Gaussia* luciferase reporter gene-bearing lentivirus pseudotyped with the wild-type S or B.1.617.2 (Delta) variant S protein in each well of a 96-well plate. Virus was incubated with 4-fold serial dilutions of serum/lung homoginates for 1 hour at 37°C (final dilutions 1:40, 1:160, 1:640, 1:2560, 1:10240, and no serum control). Media (DMEM (Gibco, 11965-092) supplemented with 10% (vol/vol) fetal bovine serum (Gibco, 11965-092) and 1% (vol/vol) penicillin/streptomycin (HyClone, SV30010)) was removed from seeded HEK293T/ACE2 cells (BEI NR-52511) and replaced with the virus/serum mixture. Cells were cultured for 6 hours at 37°C before changing to fresh media. *Gaussia* luciferase activity was measured at 24, 48, and 72 hours after media change. For luciferase measurement, 20 μL of supernatant was collected from each well and transferred to a white non-sterile 96-well plate. To each well, 20 μL of *Gaussia* luciferase substrate (0.1M Tris [MilliporeSigma, T6066] pH 7.4, 0.3M sodium ascorbate [Spectrum, S1349], 10 μM coelenterazine [GoldBio, CZ2.5]) was added, and luminescence was immediately read by a BioTek Cytation5 plate reader. 50% neutralization titers (NT_50_) were determined by least-squares-fit non-linear regression in GraphPad Prism5.

### Intravascular staining for discrimination between circulating and resident cells

Anti-CD45-PE (Clone 30-F11, BD Biosciences) (3 µg in 100 µL sterile PBS) was injected i.v. 10 minutes prior to sacrifice to label circulating lymphocytes, while resident lymphocytes are protected from labeling (Anderson et al., 2014; Wilk et al., 2017) Peripheral blood was collected at time of sacrifice and checked by flow cytometry to confirm that >90% of circulating lymphocytes were CD45-PE+.

### Tissue dissociation and flow cytometry

Lungs were isolated and processed into single cell suspension using gentleMACS tissue dissociator and mouse lung dissociation kit (Miltenyi Biotec Ref. 130-095-927) followed by 40 μm filtration and red blood cells lysis with ACK lysis buffer. Cells from each mouse were resuspended in T cell media media (RPMI 1640 supplemented with 0.1% Gentamicin antibiotic, 10% HI-FBS, Glutamax, and 5×10-5 M BME) and incubated for 4-5 hr at 37°C with protein transport inhibitor cocktail (eBioscience). Stimulation was done either nonspecifically with PMA (50 ng/ml) /Ionomycin (500 ng/ml) or specifically using two pools of Spike peptides covering the C- and N-terminal (PepTivator SARS-CoV-2 Prot_S1 and PepTivator SARS-CoV-2 Prot_S+) (Miltenyi Biotec Ref. 130-126-701& 130-126-700), Peptide pools were used at a final concentration of 1 μg/ml each peptide. Non-stimulated (DMSO only) group was also included as negative control.

Following stimulation, cells were washed with cold PBS prior to staining with LIVE/DEAD Zombie NIR Fixable Viability dye (Biolegend Cat. 423105) for 30 min at 4°C. Cells were then washed 2x with PBS supplemented with 1% HI-FBS (1% FBS) (FACS buffer) and resuspended in Fc Block (clone 93) (eBioscience Ref. 14-0161-86) at 4°C for 5 min before surface staining with a cocktail of the following antibodies for 20 min at 4°C: CD3 V450 (Clone 17A2) (BD Cat. 561389), CD4 BV750 (Clone H129.19) (BD Cat. 747275), CD44 PerCP Cy5.5 (Clone IM7) (BD Cat. 560570), CD62L BV605 (Clone MEL-14) (BD Cat. 563252), CD69 BV711 (Clone HI.2F3) (BD Cat. 740664), CD103 PE-CF594 (Clone M290) (BD Cat. 565849). After two washes in FACS buffer cells were resuspended in IC fixation buffer (eBioscience Cat. 00-8222-49) and incubated for 20 minutes at RT. Following permeabilization (eBioscience Cat. 00-8333-56), intracellular staining (30 minutes at 4°C) was done using a cocktail of the following antibodies: IFNγ FITC (clone XMG1.2) (eBioscience Cat. 11-7311-82), IL-17 PE CY7 (clone eBio17B7) (eBioscience Cat. 25-7177-82), IL-5 APC (clone TRFK5) (BD Cat. 554396). For examination of CD8+ population, the same panel was used with the replacement of CD4 BV750 antibody with CD8 APC (clone 53-6.7) (Biolegend Cat. 100712) and probing for IFNγ only. Fluorescence minus one or isotype control antibodies were used as negative controls. Finally, cells were washed with permeabilization buffer and resuspended in FACS buffer. Samples were collected on a Cytek Aurora flow cytometer (Cytekbio). Analysis was performed using FlowJo software, version 10.8.0 according to the gating strategy outlined in Figures S1 and S2. The number of cells within each population was calculated by multiplying the frequency of live singlets in the population of interest by the total number of cells in each sample.

For the analysis of T_FH_ and GC B cell populations, mediastinal lymph nodes were harvested, and single-cell suspensions were generated in processing media (Iscove’s Modified Dulbecco’s Medium (IMDM) + 4% FBS) by passing tissue through a 100μm cell strainer. Cells were incubated in 0.84% NH_4_Cl for 3 minutes to lyse red blood cells. After washing with processing media, cells were resuspended in FACS buffer (PBS + 4% FBS) supplemented with Fc block (clone 93; BioLegend Cat. 101320) and incubated at 4°C for 5 minutes before staining. For surface staining, cells were incubated for 30 minutes at 4°C with the following antibodies, as indicated: Ghost viability dye (1:400; Tonbo Biosciences Cat. 13-0870-T100); CD4-AF488 (1:300; clone GK1.5; R&D Systems Cat. FAB554G), CD44-BV421 (1:300; clone IM7; BD Biosciences Cat. 563970); CD62L-APC-eFluor780 (1:300; clone MEL-14; ThermoFisher Cat. 47-0621-82); PD-1-PE-Cy7 (1:50; clone 29F.1A12; BioLegend Cat. 135216); Cxcr5-APC (1:50; clone SPRCL5; ThermoFisher Cat. 17-7185-82); CD19:APC (1:200; clone: GL7, BioLegend Cat. 144610); GL-7: PerCP-Cy5.5 (1:200; clone: 1D3, BioLegend Cat.152410); Fas-BV421 (1:300; clone Jo2; BD Biosciences Cat. 562633). Cells were then washed 2X with FACS buffer and fixed using the eBioscience Foxp3 transcription factor staining kit (ThermoFisher Cat. 00-5523-00), according to the manufacturer’s instructions. Following fixation, cells were washed 1X with eBioscience 1X permeabilization buffer, and 2X with FACS buffer before resuspension in FACS buffer for analysis. Samples were analyzed on a BD FACS Canto II, and full analysis was performed using FlowJo software, version 10.8.0.

### Splenocyte stimulation and cytokine ELISA assays

Following dissociation and red blood cell lysis using ACK buffer, a single-cell suspension was plated at 2.5 × 10^6^ cells/well of complete T cell medium (RPMI, 10% FBS, 10 μg/ml gentamicin, 5 × 10^−5^ M 2-mercaptoethanol) and stimulated with 1 μg/ml S protein or with medium alone as a negative control. The supernatant was collected on day 3 post stimulation. The production of IFNγ (R&D Cat. DY485-05), IL-5 (Invitrogen Cat. 88-7054-88) and IL-17 (Invitrogen Cat. 88-7371-88) was quantified by a sandwich ELISA according to the manufacturer’s instructions. Plates were read at A_450_ on a SpectraMax i3x® plate reader and concentration calculated based on the standard curve.

### ELISA for S specific Igs

Corning Costar high binding 96 well ELISA plates (Ref. 9018) were coated with 1µg/ml of S protein in 1X PBS at 4°C overnight. Plates were washed 3x with PBS-Tween 20, blocked for 2 hr at RT with Elisa diluent (Invitrogen) (Ref. 00-4202-56), then washed 2x with PBS-Tween 20. Serial dilutions of serum (1:500, 1:2500 and 1:12500 for IgG and IgA) and lung supernatant (1:500, 1:2500 and 1:12500 for IgG, 1:50 and 1:250 for IgA) samples were added, and incubated for 2 hr at RT. Plates were washed 4x with PBS-Tween 20 and incubated with secondary anti-mouse IgG HRP (Abcam Ref. ab6789) or anti-mouse IgA HRP antibodies for 1 hr at RT. Plates were developed for 5-15 min at RT using 100µl of TMB (Invitrogen Ref. 50-112-9758) and the reaction stopped with H2SO4. Plates were read at A_450_ on a SpectraMax i3x® plate reader. To evaluate antibody avidity, sodium thiocyanate (0-3M) was added for 20 min at 37°C, prior to blocking and addition of detection antibody (Macdonald et al., 1988). IgG RAI was calculated: (A450 at 2M) / (A450 at 0 M) *100, and IgA RAI was calculated by (A450 at 1M) / (A450 at 0 M) *100.

To determine IgG isotypes of S-specific antibodies, plates were coated, blocked and incubated with serum samples (1:5000), or lung homogenates (1:1000) as described above. Rat anti mouse antibodies specific for IgG subtypes (IgG1, IgG2b, IgG2C& IgG3-Invitrogen) (Ref. 88-50630-88 & 88-50670-22) were added (1:1000 dilution) and incubated for 1 hr at RT. Plates were washed 4x with PBS-Tween 20, incubated with secondary anti-Rat IgG HRP (1:5000, Rockland Ref. 612-1302) for 1 hour at RT, then developed with TMB as above.

### Lung Histopathology and Immunohistochemistry

Lung samples from mice were processed per a standard protocol. Briefly, the tissues were fixed in 10% neutral buffered formalin. Tissues were processed and embedded in paraffin. Five-micrometer sections (3 per tissue) were stained with hematoxylin and eosin by the Comparative Pathology & Mouse Phenotyping Shared Resource at The Ohio State University. A board-certified veterinary pathologist (K.N.C.) was blind to the experimental groups, and sections were scored qualitatively on a scale ranging from 0 to 5 for the degree of cellularity and consolidation, the thickness of the alveolar walls, degeneration and necrosis, edema, hemorrhage, infiltrating alveolar/interstitial polymorphonuclear cells (PMNs), intrabronchial PMNs, perivascular and peribronchial lymphocytes and plasma cells, and alveolar macrophages. The total inflammation score was calculated by totaling the qualitative assessments in each category.

Immunohistochemistry to detect nucleocapsid protein expression in lung sections was conducted by HistoWiz, Inc. Briefly, tissue sections were stained with a rabbit monoclonal anti-nucleocapsid antibody (Genetex # GTX635686) using standard methodology.

### SARS-CoV-2 MA10 infection and viral titer measurements

Anesthetized (isoflourane) mice were intranasaly infected with 10^5^ PFU of SARS-CoV-2 MA10 diluted in PBS where indicated. Clinical signs of disease (weight loss) were monitored daily. Mice were euthanized by isoflurane overdose at 2 days post infection and samples for titer (caudal right lung lobe and nasal septum) and histopathological analyses (left lung lobe) were collected. Importantly, mice were randomized and assigned to specific harvest days before the start of the experiment. Lung viral titers were determined by plaque assay. Briefly, right caudal lung lobes were homogenized in 1mL PBS using glass beads and serial dilutions of the clarified lung homogenates were added to a monolayer of Vero E6 cells. After three days CPE was examined via staining for viral nucleoprotein (Sino Biological Cat. 40143-MM08-100). The left lung lobe was stored in 10% phosphate buffered formalin for 7 days prior to removal from the BSL3 for processing. After paraffin embedding, sectioning and staining histopathological scoring was performed.

### Statistical Analysis

Serological responses, T and B cell readouts, mouse weights and viral titer data were analyzed using Graphpad Prism by the methods outlined in each figure legend.

## ACKNOWLEDGEMENTS

This work was supported by The Ohio State University COVID-19 Seed Fund and R21AI151867 (PD), R01AI125560 (PD and RD), private donor funds (S-LL), R01AI090060 and RM1HG008935 COVID-19 supplement award (JL), R01AI130110, R01HL154001, R01CA260582, and an American Lung Association COVID-19 and Emerging Respiratory Viruses Research Award (JSY), U19AI131386-04S1 (MEP), Nationwide Children’s Hospital COVID-19 Seed Fund (MEP), R01AI134972 (KJO). The Comparative Pathology & Digital Imaging Shared Resource and KNC are supported in part by P30CA16058MS. MMS is supported by a doctoral fellowship from the Egyptian Bureau of Higher Education. AZ is supported by an NSF-GRFP fellowship. KAR is supported by The Ohio State University College of Medicine Advancing Research in Infection and Immunity Fellowship Program. JMB was supported by T32GM068412. We thank Jason McLellan for the plasmid expressing the HexaPro stabilized version of the S protein.

## AUTHOR CONTRIBUTIONS

MMS, KC, JH, JMB, KAR, conducted immunological analysis, MM, AZ, AK conducted SARS-CoV-2 challenge, euthanasia, and TCID50 assays, JE and CZ conducted virus neutralization assays, KCM and ML purified MA10 virus, SC produced S protein, KNC conducted histopathological analysis. PD designed the studies and PD and MMS analyzed data and wrote the manuscript. JSY supervised BSL-3 work, JL and MEP supervised production of MA10 virus, MEP supervised production of S protein, S-LL supervised virus neutralization assays, KJO supervised Tfh and GC B cell analysis, RD and PD provided BcfA.

---

**Figure S1:**
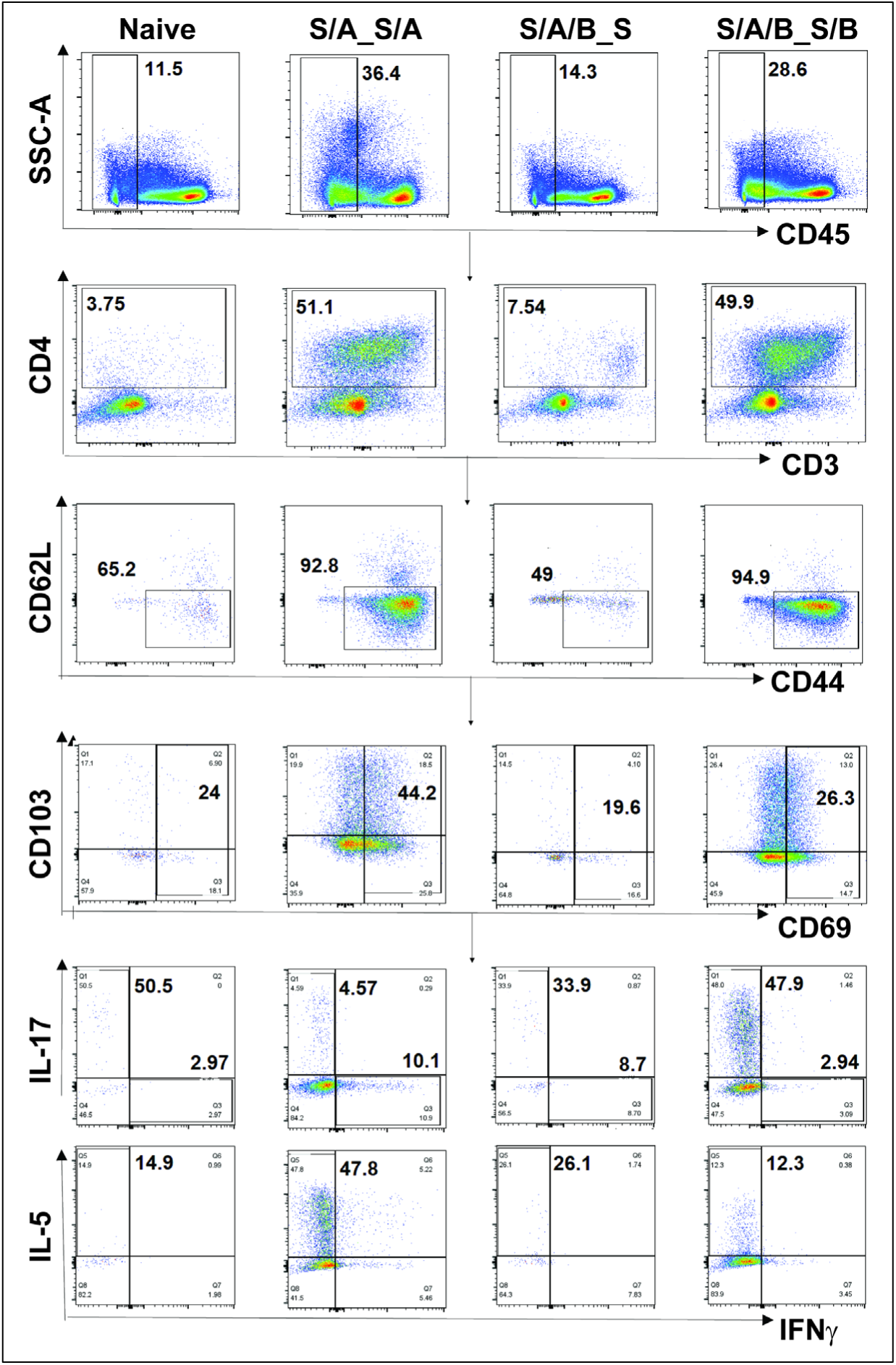
Gating strategy to identify resident and circulating memory T cells in the lungs. Mice were injected with CD45 PE antibody 10 minutes before sac to distinguish between resident (CD45-) and circulating (CD45+) T cells. PMA/Ionomycin stimulated cells stained with surface markers to identify antigen experienced CD4+CD44+CD62L-CD69+ CD4+ cells were fixed, pearmeabilized and followed by intracellular cytokine staining (ICS) to analyze the production of IFNγ, IL-5 and IL-17. Shown is the sequential gating strategy to identify lymphocytes by forward and side scatter, single cells, live cells, CD45-cells, CD3+CD4+ cells, CD44+CD62L-cells and CD69+ cells. The proportion of IFNγ+, IL-17+ and IL-5+ cells within the CD3+CD4+CD45-CD44+CD62L-CD69+ fraction was determined. The same sequential gating strategy was used to identify CD45+ cells.

**Figure S2:**
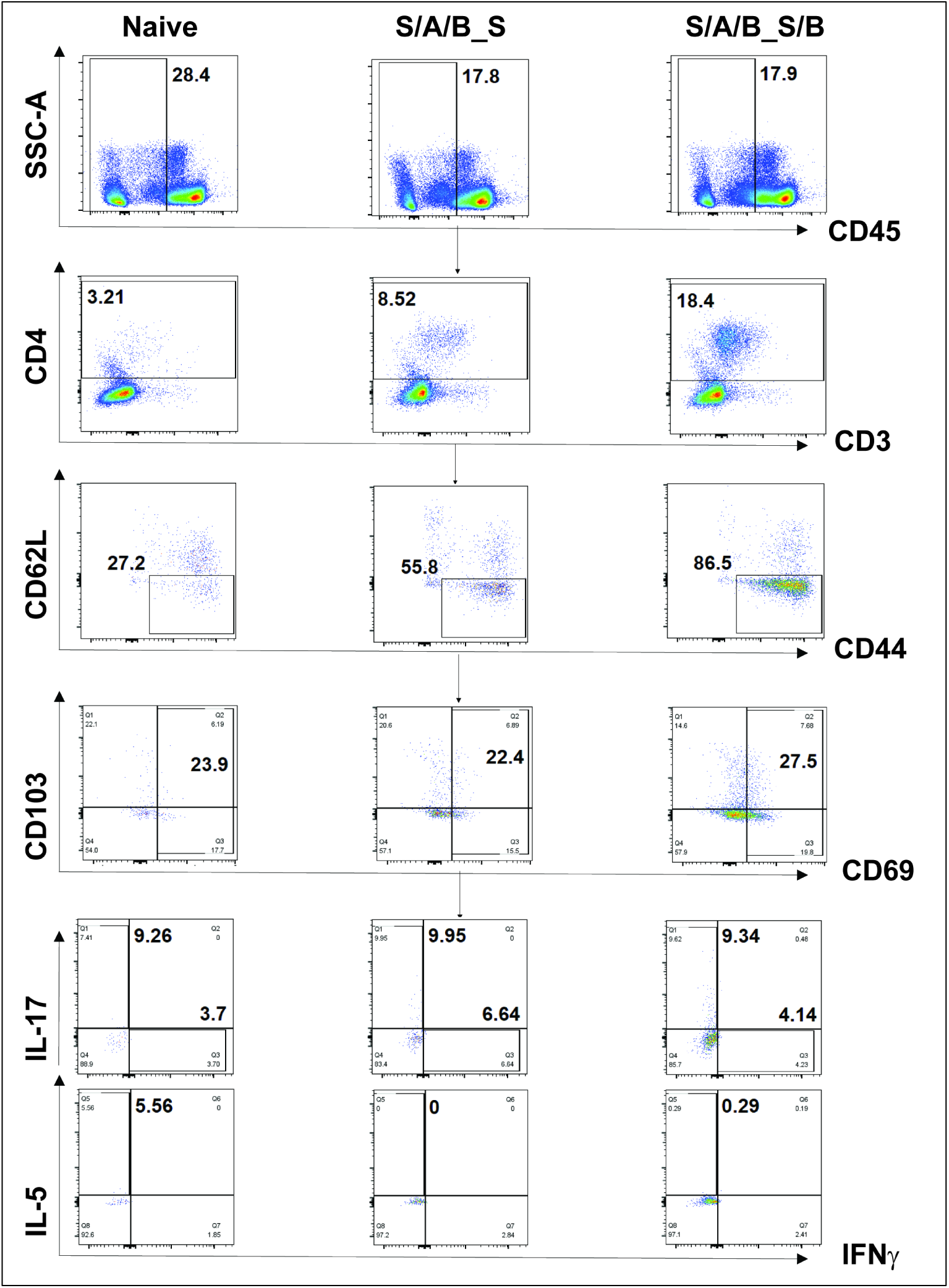
Gating strategy to identify antigen-specific resident and circulating memory T cells in the lungs. Dissociated lung cells stimulated with S1 and S2 peptide pools were stained and sequentially gated to identify the CD45- and CD45+ fractions, as described in Figure S1.

**Figure S3:**
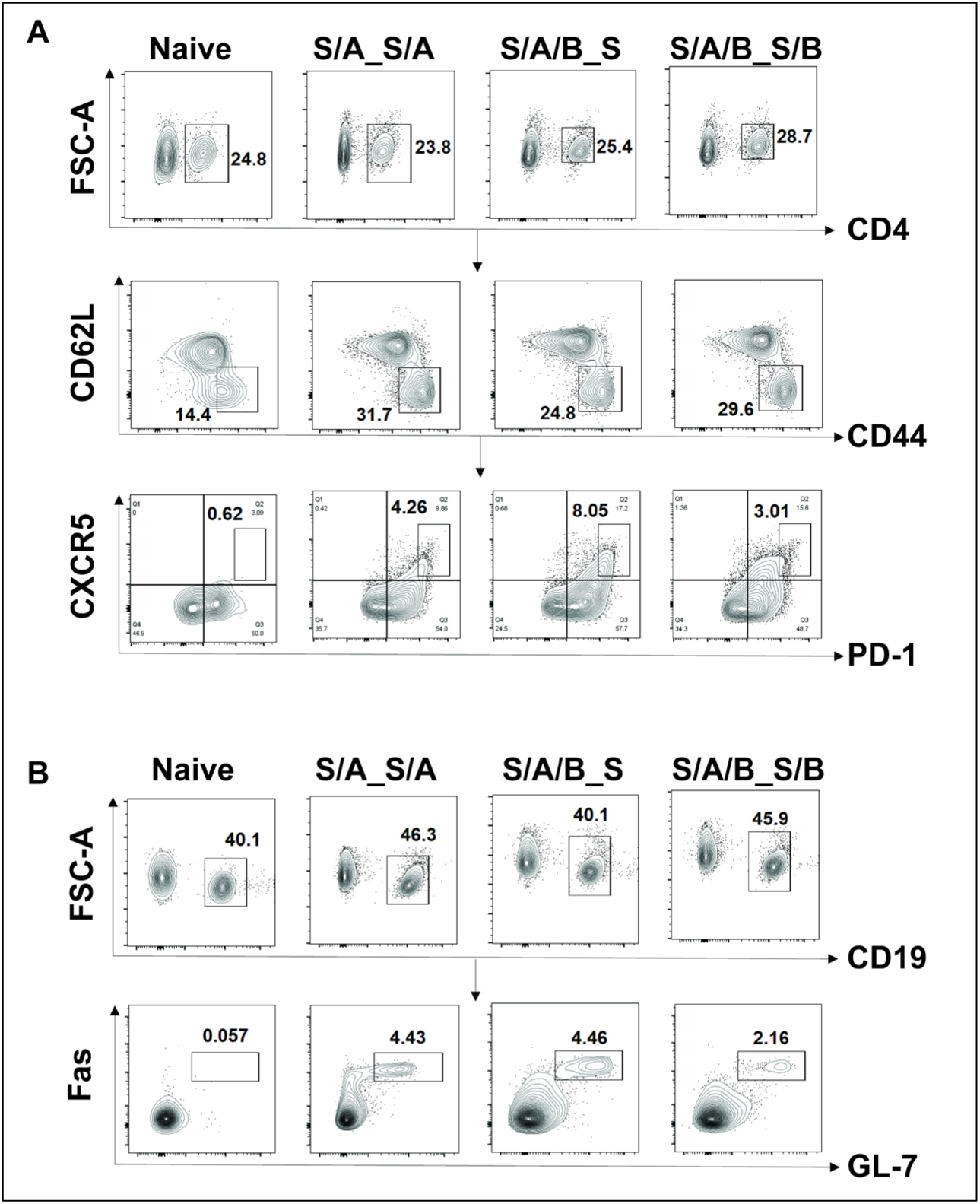
Gating strategy to identify PD-1^hi^ CXCR5^hi^ Tfh cells and Fas+GL7+ GC B cells in the draining mediastinal lymph node. (A) Dissociated lymph node cells stained with antibodies against CD4, CD44, CD62L, PD-1 and CXCR5 to identify CD44^+^PD-1^high^ CXCR5^hi^gh Tfh cells. (B) Lymph node cells stained with antibodies against GC B cells (Fas+GL7+).

**Figure S4:**
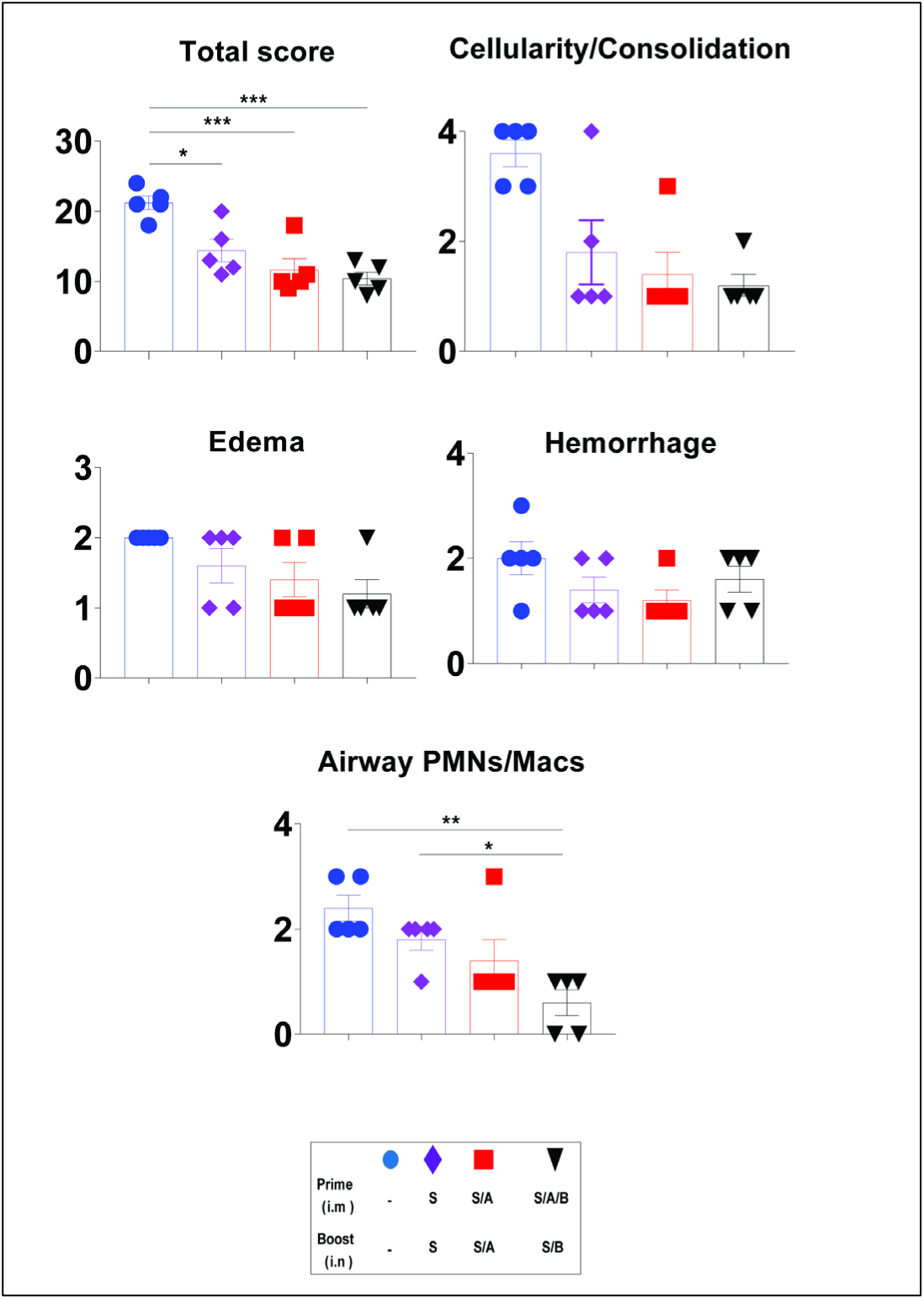
Semi-quantitative histopathological scoring of lung sections from MA10 challenged C57BL/6 mice. Shown is the total score, cellularity/consolidation, edema, hemorrhage, and infiltration of PMNs and macrophages in the airways in the lungs of naïve challenged mice and mice immunized with S alone, S/A or S/A/B_S/B.

**Figure S5:**
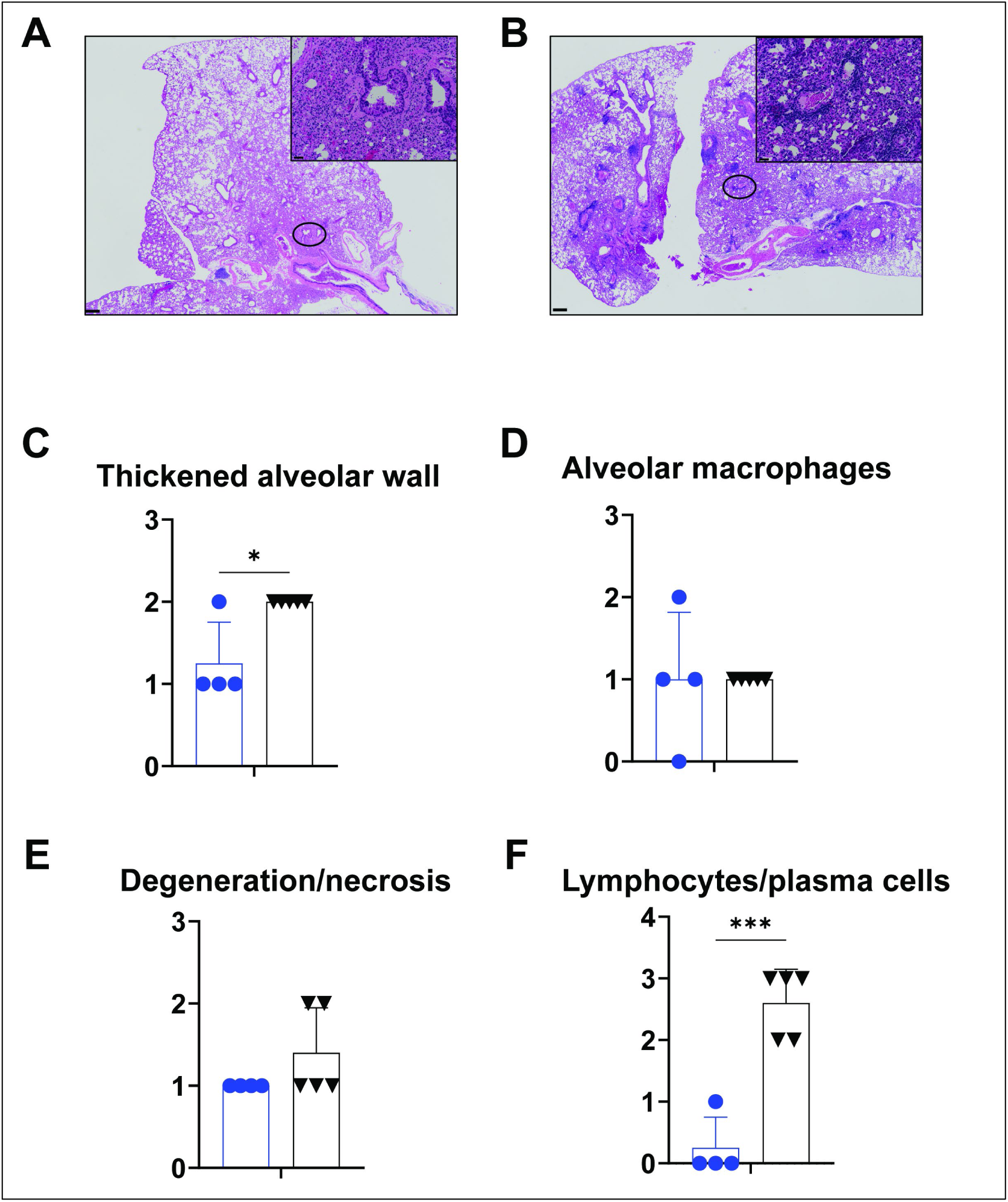
IL-17 KO mice immunized with S/A/B_S/B display lung pathology after challenge with SARS-CoV-2 MA10. Formalin fixed, paraffin embedded lungs harvested on d3 post-infection were sectioned and stained with H&E to evaluate inflammation and tissue damage. (A) Unimmunized IL-17 KO mice, (B) S/A/B_S/B immunized mice. Semi-quantitative scoring of (C) alveolar wall thickness (D) presence of alveolar macrophages (E) degeneration and necrosis and (F) presence of lymphocytes and plasma cells is shown 4-5 samples/group, evaluated by Student’s t-test. *, p< 0.05, ***, p<0.001.

**Table S1.**
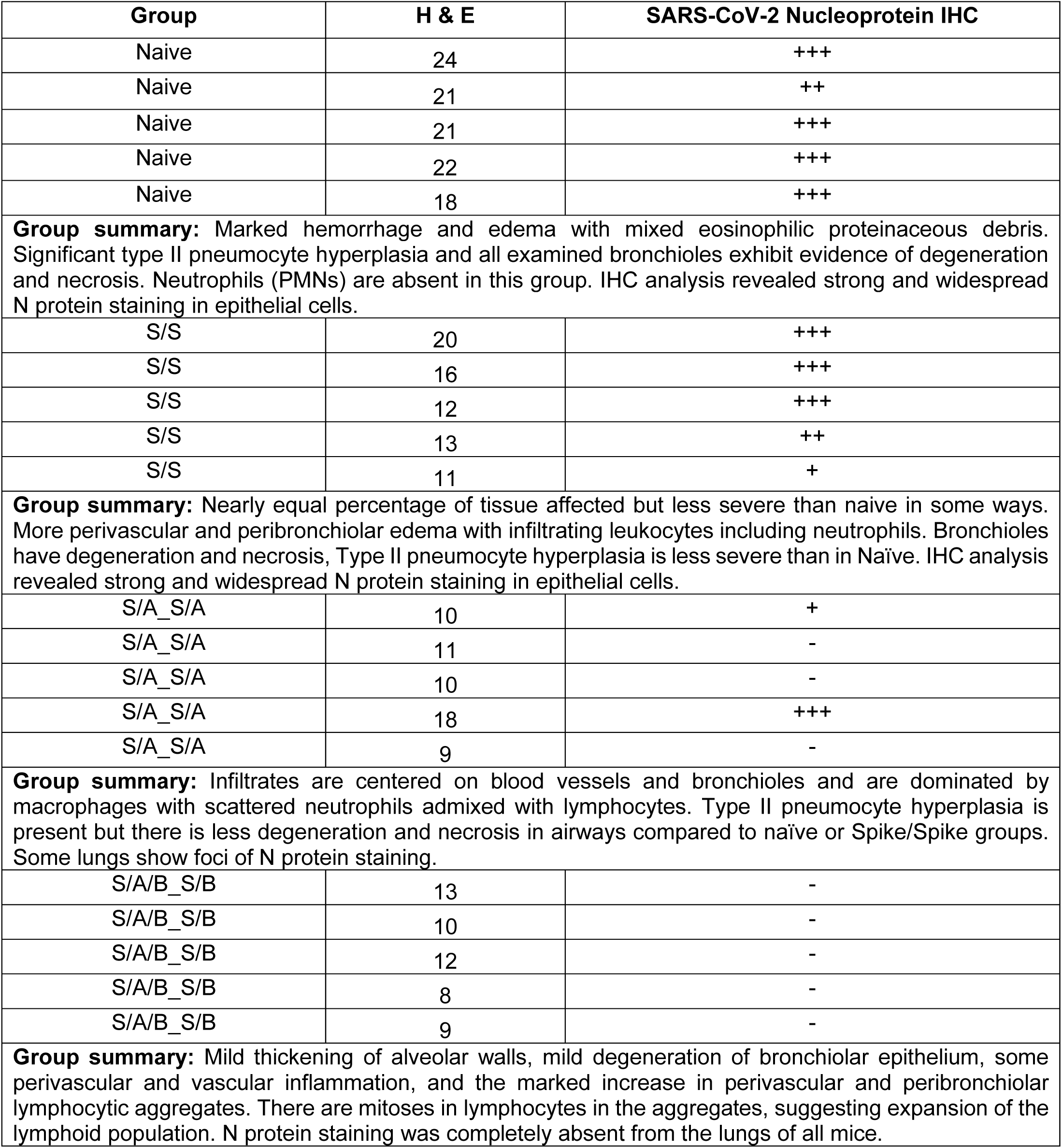
Day 2 histopathological scoring in the lungs of MA10 challenged mice

Day 2 histopathological scoring in the lungs of MA10 challenged mice
| Group | H & E | SARS-CoV-2 Nucleoprotein IHC |
| --- | --- | --- |
| Naive | 24 | +++ |
| Naive | 21 | ++ |
| Naive | 21 | +++ |
| Naive | 22 | +++ |
| Naive | 18 | +++ |
| **Group summary:** Marked hemorrhage and edema with mixed eosinophilic proteinaceous debris. Significant type II pneumocyte hyperplasia and all examined bronchioles exhibit evidence of degeneration and necrosis. Neutrophils (PMNs) are absent in this group. IHC analysis revealed strong and widespread N protein staining in epithelial cells. |  |  |
| S/S | 20 | +++ |
| S/S | 16 | +++ |
| S/S | 12 | +++ |
| S/S | 13 | ++ |
| S/S | 11 | + |
| **Group summary:** Nearly equal percentage of tissue affected but less severe than naive in some ways. More perivascular and peribronchiolar edema with infiltrating leukocytes including neutrophils. Bronchioles have degeneration and necrosis, Type II pneumocyte hyperplasia is less severe than in Naïve. IHC analysis revealed strong and widespread N protein staining in epithelial cells. |  |  |
| S/A_S/A | 10 | + |
| S/A_S/A | 11 | - |
| S/A_S/A | 10 | - |
| S/A_S/A | 18 | +++ |
| S/A_S/A | 9 | - |
| **Group summary:** Infiltrates are centered on blood vessels and bronchioles and are dominated by macrophages with scattered neutrophils admixed with lymphocytes. Type II pneumocyte hyperplasia is present but there is less degeneration and necrosis in airways compared to naïve or Spike/Spike groups. Some lungs show foci of N protein staining. |  |  |
| S/A/B_S/B | 13 | - |
| S/A/B_S/B | 10 | - |
| S/A/B_S/B | 12 | - |
| S/A/B_S/B | 8 | - |
| S/A/B_S/B | 9 | - |
| **Group summary:** Mild thickening of alveolar walls, mild degeneration of bronchiolar epithelium, some perivascular and vascular inflammation, and the marked increase in perivascular and peribronchiolar lymphocytic aggregates. There are mitoses in lymphocytes in the aggregates, suggesting expansion of the lymphoid population. N protein staining was completely absent from the lungs of all mice. |  |  |

## References

1. Agrawal, A.S.,, Tao, X.,, Algaissi, A.,, Garron, T.,, Narayanan, K.,, et al. (2016). Immunization with inactivated Middle East Respiratory Syndrome coronavirus vaccine leads to lung immunopathology on challenge with live virus. 12, 2351–2356.

2. Allen, A.C., Wilk, M.M., Misiak, A., Borkner, L., Murphy, D., and Mills, K.H.G. (2018). Sustained protective immunity against Bordetella pertussis nasal colonization by intranasal immunization with a vaccine-adjuvant combination that induces IL-17-secreting TRM cells. Mucosal Immunol 11, 1763–1776.

3. Amanat, F., and Krammer, F. (2020). SARS-CoV-2 Vaccines: Status Report. Immunity 52, 583–589.

4. Anderson, K.G., Mayer-Barber, K., Sung, H., Beura, L., James, B.R., Taylor, J.J., Qunaj, L., Griffith, T.S., Vezys, V., Barber, D.L., et al. (2014). Intravascular staining for discrimination of vascular and tissue leukocytes. Nat Protoc 9, 209–222.

5. Arunachalam, P.S., Walls, A.C., Golden, N., Atyeo, C., Fischinger, S., Li, C., Aye, P., Navarro, M.J., Lai, L., Edara, V.V., et al. (2021). Adjuvanting a subunit COVID-19 vaccine to induce protective immunity. Nature 594, 253–258.

6. Bai, Y., Yao, L., Wei, T., Tian, F., Jin, D.Y., Chen, L., and Wang, M. (2020). Presumed Asymptomatic Carrier Transmission of COVID-19. JAMA 323, 1406–1407.

7. Bolles, M.,, Deming, D.,, Long, K.,, Agnihothram, S.,, Whitmore, A.,, et al. (2011). A double-inactivated severe acute respiratory syndrome coronavirus vaccine provides incomplete protection in mice and induces increased eosinophilic proinflammatory pulmonary response upon challenge. 85, 12201–12215.

8. Case, J.B.,, Winkler, E.S.,, Errico, J.M.,, Diamond, M.S., and (2021). On the road to ending the COVID-19 pandemic: Are we there yet? 557, 70.

9. Crotty, S. (2014). T follicular helper cell differentiation, function, and roles in disease. Immunity 41, 529–542.

10. Cyster, J.G., and Allen, C.D.C. (2019). B Cell Responses: Cell Interaction Dynamics and Decisions. Cell 177, 524–540.

11. Czub, M.,, Weingartl, H.,, Czub, S.,, He, R.,, Cao, J., and (2005). Evaluation of modified vaccinia virus Ankara based recombinant SARS vaccine in ferrets. 23, 2273–2279.

12. DiPiazza, A.T., Leist, S.R., Abiona, O.M., Moliva, J.I., Werner, A., Minai, M., Nagata, B.M., Bock, K.W., Phung, E., Schafer, A., et al. (2021). COVID-19 vaccine mRNA-1273 elicits a protective immune profile in mice that is not associated with vaccine-enhanced disease upon SARS-CoV-2 challenge. Immunity 54, 1869–1882 e1866.

13. Fulginiti, V.A.,, Eller, J.J.,, Downie, A.W.,, Kempe, C.H., and (1967). Altered Reactivity to Measles Virus: Atypical Measles in Children Previously Immunized With Inactivated Measles Virus Vaccines. 202, 1075–1080.

14. Gandhi, M., Yokoe, D.S., and Havlir, D.V. (2020). Asymptomatic Transmission, the Achilles’ Heel of Current Strategies to Control Covid-19. N Engl J Med 382, 2158–2160.

15. Graham, B.S., Henderson, G.S., Tang, Y.W., Lu, X., Neuzil, K.M., and Colley, D.G. (1993). Priming immunization determines T helper cytokine mRNA expression patterns in lungs of mice challenged with respiratory syncytial virus. J Immunol 151, 2032–2040.

16. Grifoni, A., Weiskopf, D., Ramirez, S.I., Mateus, J., Dan, J.M., Moderbacher, C.R., Rawlings, S.A., Sutherland, A., Premkumar, L., Jadi, R.S., et al. (2020). Targets of T Cell Responses to SARS-CoV-2 Coronavirus in Humans with COVID-19 Disease and Unexposed Individuals. Cell 181, 1489–1501 e1415.

17. Halstead, S.B., and Katzelnick, L. (2020). COVID-19 Vaccines: Should We Fear ADE? J Infect Dis 222, 1946–1950.

18. Hellerstein, M. (2020). What are the roles of antibodies versus a durable, high quality T-cell response in protective immunity against SARS-CoV-2? Vaccine X 6, 100076.

19. Honda-Okubo, Y.,, Barnard, D.,, Ong, C.H.,, Peng, B.-H.,, Tseng, C.-T.K.,, et al. (2015). Severe acute respiratory syndrome-associated coronavirus vaccines formulated with delta inulin adjuvants provide enhanced protection while ameliorating lung eosinophilic immunopathology. 89, 2995–3007.

20. Hotez, P.J., Bottazzi, M.E., and Corry, D.B. (2020). The potential role of Th17 immune responses in coronavirus immunopathology and vaccine-induced immune enhancement. Microbes Infect 22, 165–167.

21. Hsieh, C.L., Goldsmith, J.A., Schaub, J.M., DiVenere, A.M., Kuo, H.C., Javanmardi, K., Le, K.C., Wrapp, D., Lee, A.G., Liu, Y., et al. (2020). Structure-based design of prefusion-stabilized SARS-CoV-2 spikes. Science 369, 1501–1505.

22. Jennings-Gee, J., Quataert, S., Ganguly, T., D’Agostino, R., Jr., Deora, R., and Dubey, P. (2018). The Adjuvant Bordetella Colonization Factor A Attenuates Alum-Induced Th2 Responses and Enhances Bordetella pertussis Clearance from Mouse Lungs. Infect Immun 86.

23. Kaneko, N., Kuo, H.H., Boucau, J., Farmer, J.R., Allard-Chamard, H., Mahajan, V.S., Piechocka-Trocha, A., Lefteri, K., Osborn, M., Bals, J., et al. (2020). Loss of Bcl-6-Expressing T Follicular Helper Cells and Germinal Centers in COVID-19. Cell 183, 143–157 e113.

24. Kared, H., Redd, A.D., Bloch, E.M., Bonny, T.S., Sumatoh, H., Kairi, F., Carbajo, D., Abel, B., Newell, E.W., Bettinotti, M.P., et al. (2021). SARS-CoV-2-specific CD8+ T cell responses in convalescent COVID-19 individuals. J Clin Invest 131.

25. Ketas, T.J.,, Chaturbhuj, D.,, Cruz Portillo, V.M.,, Francomano, E.,, Golden, E.,, et al. (2021). Antibody responses to SARS-CoV-2 mrna vaccines are detectable in Saliva. 6, 116–134.

26. Kim, H.W., Canchola, J.G., Brandt, C.D., Pyles, G., Chanock, R.M., Jensen, K., and Parrott, R.H. (1969). Respiratory syncytial virus disease in infants despite prior administration of antigenic inactivated vaccine. Am J Epidemiol 89, 422–434.

27. Krautler, N.J., Suan, D., Butt, D., Bourne, K., Hermes, J.R., Chan, T.D., Sundling, C., Kaplan, W., Schofield, P., Jackson, J., et al. (2017). Differentiation of germinal center B cells into plasma cells is initiated by high-affinity antigen and completed by Tfh cells. J Exp Med 214, 1259–1267.

28. Lavelle, E.C., and Ward, R.W. (2021). Mucosal vaccines - fortifying the frontiers. Nat Rev Immunol.

29. Le Bert, N., Tan, A.T., Kunasegaran, K., Tham, C.Y.L., Hafezi, M., Chia, A., Chng, M.H.Y., Lin, M., Tan, N., Linster, M., et al. (2020). SARS-CoV-2-specific T cell immunity in cases of COVID-19 and SARS, and uninfected controls. Nature 584, 457–462.

30. Leist, S.R., Dinnon, K.H., 3rd, Schafer, A., Tse, L.V., Okuda, K., Hou, Y.J., West, A., Edwards, C.E., Sanders, W., Fritch, E.J., et al. (2020). A Mouse-Adapted SARS-CoV-2 Induces Acute Lung Injury and Mortality in Standard Laboratory Mice. Cell 183, 1070–1085 e1012.

31. Liang, Z., Zhu, H., Wang, X., Jing, B., Li, Z., Xia, X., Sun, H., Yang, Y., Zhang, W., Shi, L., et al. (2020). Adjuvants for Coronavirus Vaccines. Front Immunol 11, 589833.

32. Lippi, G., Sanchis-Gomar, F., and Henry, B.M. (2020). COVID-19: unravelling the clinical progression of nature’s virtually perfect biological weapon. Ann Transl Med 8, 693.

33. Lycke, N. (2012). Recent progress in mucosal vaccine development: potential and limitations. Nat Rev Immunol 12, 592–605.

34. Macdonald, R.A., Hosking, C.S., and Jones, C.L. (1988). The measurement of relative antibody affinity by ELISA using thiocyanate elution. J Immunol Methods 106, 191–194.

35. Mades, A.,, Chellamathu, P.,, Lopez, L.,, Kojima, N.,, MacMullan, M.A.,, et al. (2021). Detection of persistent SARS-CoV-2 IgG antibodies in oral mucosal fluid and upper respiratory tract specimens following COVID-19 mRNA vaccination. 2021.2005.2006.21256403.

36. Matuchansky, C. (2021). Mucosal immunity to SARS-CoV-2: a clinically relevant key to deciphering natural and vaccine-induced defences. Clin Microbiol Infect.

37. McPherson, C.,, Chubet, R.,, Holtz, K.,, Honda-Okubo, Y.,, Barnard, D.,, et al. (2016). Development of a SARS Coronavirus Vaccine from Recombinant Spike Protein Plus Delta Inulin Adjuvant. 1403, 269–284.

38. Nader, P.R.,, Horwitz, M.S.,, Rousseau, J., and (1968). Atypical exanthem following exposure to natural measles: Eleven cases in children previously inoculated with killed vaccine. 72, 22–28.

39. Orlov, M.,, Wander, P.L.,, Morrell, E.D.,, Mikacenic, C.,, Wurfel, M.M., and (2020). A Case for Targeting Th17 Cells and IL-17A in SARS-CoV-2 Infections. 205, 892–898.

40. Pacha, O.,, Sallman, M.A.,, Evans, S.E., and (2020). COVID-19: a case for inhibiting IL-17? 20, 345–346.

41. Parackova, Z.,, Bloomfield, M.,, Klocperk, A.,, Sediva, A., and (2021). Neutrophils mediate Th17 promotion in COVID-19 patients. 109, 73–76.

42. Peng, Y., Mentzer, A.J., Liu, G., Yao, X., Yin, Z., Dong, D., Dejnirattisai, W., Rostron, T., Supasa, P., Liu, C., et al. (2020). Broad and strong memory CD4(+) and CD8(+) T cells induced by SARS-CoV-2 in UK convalescent individuals following COVID-19. Nat Immunol 21, 1336–1345.

43. Roces, C.B., Hussain, M.T., Schmidt, S.T., Christensen, D., and Perrie, Y. (2019). Investigating Prime-Pull Vaccination through a Combination of Parenteral Vaccination and Intranasal Boosting. Vaccines (Basel) 8.

44. Russell, M.W., Moldoveanu, Z., Ogra, P.L., and Mestecky, J. (2020). Mucosal Immunity in COVID-19: A Neglected but Critical Aspect of SARS-CoV-2 Infection. Front Immunol 11, 611337.

45. S, G.B.,, S, H.G.,, W, T.Y.,, X, L.,, M, N.K.,, et al. (1993). Priming immunization determines T helper cytokine mRNA expression patterns in lungs of mice challenged with respiratory syncytial virus - PubMed. 151, 2032–2040.

46. Sarmiento-Monroy, J.C.,, Parra-Medina, R.,, Garavito, E.,, Rojas-Villarraga, A., and (2021). T Helper 17 Response to Severe Acute Respiratory Syndrome Coronavirus 2: A Type of Immune Response with Possible Therapeutic Implications. 34, 190–200.

47. See, R.H.,, Zakhartchouk, A.N.,, Petric, M.,, Lawrence, D.J.,, Mok, C.P.Y.,, et al. (2006). Comparative evaluation of two severe acute respiratory syndrome (SARS) vaccine candidates in mice challenged with SARS coronavirus. 87, 641–650.

48. Seephetdee, C., Buasri, N., Bhukhai, K., Srisanga, K., Manopwisedjaroen, S., Lertjintanakit, S., Phueakphud, N., Pakiranay, C., Kangwanrangsan, N., Srichatrapimuk, S., et al. (2021). Mice Immunized with the Vaccine Candidate HexaPro Spike Produce Neutralizing Antibodies against SARS-CoV-2. Vaccines (Basel) 9.

49. Sekine, T., Perez-Potti, A., Rivera-Ballesteros, O., Stralin, K., Gorin, J.B., Olsson, A., Llewellyn-Lacey, S., Kamal, H., Bogdanovic, G., Muschiol, S., et al. (2020). Robust T Cell Immunity in Convalescent Individuals with Asymptomatic or Mild COVID-19. Cell 183, 158–168 e114.

50. Sui, Y., Li, J., Zhang, R., Prabhu, S.K., Andersen, H., Venzon, D., Cook, A., Brown, R., Teow, E., Velasco, J., et al. (2021). Protection against SARS-CoV-2 infection by a mucosal vaccine in rhesus macaques. JCI Insight 6.

51. Sukumar, N., Love, C.F., Conover, M.S., Kock, N.D., Dubey, P., and Deora, R. (2009). Active and passive immunizations with Bordetella colonization factor A protect mice against respiratory challenge with Bordetella bronchiseptica. Infect Immun 77, 885–895.

52. Tan, Y., Liu, F., Xu, X., Ling, Y., Huang, W., Zhu, Z., Guo, M., Lin, Y., Fu, Z., Liang, D., et al. (2020). Durability of neutralizing antibodies and T-cell response post SARS-CoV-2 infection. Front Med 14, 746–751.

53. Tostanoski, L.H., Gralinski, L.E., Martinez, D.R., Schaefer, A., Mahrokhian, S.H., Li, Z., Nampanya, F., Wan, H., Yu, J., Chang, A., et al. (2021). Protective Efficacy of Rhesus Adenovirus COVID-19 Vaccines against Mouse-Adapted SARS-CoV-2. J Virol 95, e0097421.

54. Tseng, C.T.,, Sbrana, E.,, Iwata-Yoshikawa, N.,, Newman, P.C.,, Garron, T.,, et al. (2012). Immunization with SARS coronavirus vaccines leads to pulmonary immunopathology on challenge with the SARS virus. 7.

55. Wajnberg, A., Amanat, F., Firpo, A., Altman, D.R., Bailey, M.J., Mansour, M., McMahon, M., Meade, P., Mendu, D.R., Muellers, K., et al. (2020). Robust neutralizing antibodies to SARS-CoV-2 infection persist for months. Science 370, 1227–1230.

56. Wilk, M.M., and Mills, K.H.G. (2018). CD4 TRM Cells Following Infection and Immunization: Implications for More Effective Vaccine Design. Front Immunol 9, 1860.

57. Wilk, M.M., Misiak, A., McManus, R.M., Allen, A.C., Lynch, M.A., and Mills, K.H.G. (2017). Lung CD4 Tissue-Resident Memory T Cells Mediate Adaptive Immunity Induced by Previous Infection of Mice with Bordetella pertussis. J Immunol 199, 233–243.

58. Wu, D.,, Yang, X.O., and (2020). TH17 responses in cytokine storm of COVID-19: An emerging target of JAK2 inhibitor Fedratinib. 53, 368.

59. Xu, Z.,, Shi, L.,, Wang, Y.,, Zhang, J.,, Huang, L.,, et al. (2020). Pathological findings of COVID-19 associated with acute respiratory distress syndrome. 8, 420.

60. Yasui, F.,, Kai, C.,, Kitabatake, M.,, Inoue, S.,, Yoneda, M.,, et al. (2008). Prior Immunization with Severe Acute Respiratory Syndrome (SARS)-Associated Coronavirus (SARS-CoV) Nucleocapsid Protein Causes Severe Pneumonia in Mice Infected with SARS-CoV. 181, 6337–6348.

61. Zeng, C., Evans, J.P., Pearson, R., Qu, P., Zheng, Y.M., Robinson, R.T., Hall-Stoodley, L., Yount, J., Pannu, S., Mallampalli, R.K., et al. (2020). Neutralizing antibody against SARS-CoV-2 spike in COVID-19 patients, health care workers, and convalescent plasma donors. JCI Insight 5.

62. Zhu, N.,, Zhang, D.,, Wang, W.,, Li, X.,, Yang, B.,, et al. (2020). A Novel Coronavirus from Patients with Pneumonia in China, 2019. 382, 727–733.

